# *BjuWRR1*, a CC-NB-LRR gene identified in *Brassica juncea*, confers resistance to white rust caused by *Albugo Candida*

**DOI:** 10.1101/509851

**Authors:** Heena Arora, K. Lakshmi Padmaja, Kumar Paritosh, Nitika Mukhi, A. K. Tewari, Arundhati Mukhopadhyay, Vibha Gupta, Akshay K Pradhan, Deepak Pental

## Abstract

White rust caused by oomycete pathogen *Albugo candida* is a significant disease of crucifer crops including *Brassica juncea* (mustard), a major oilseed crop of the Indian subcontinent. Earlier a resistance-conferring locus named AcB1-A5.1 was mapped in an east European gene pool line of *B. juncea* – Donskaja-IV. This line was tested along with some other lines of *B. juncea* (AABB), *B. rapa* (AA) and *B. nigra* (BB) for resistance to six isolates of *A. candida* collected from different mustard growing regions of India. Donskaja-IV was found to be completely resistant to all the tested isolates. Sequencing of a BAC spanning the locus AcB1-A5.1 showed the presence of a single CC-NB-LRR protein encoding R gene. The genomic sequence of the putative R gene with its native promoter and terminator was used for the genetic transformation of a susceptible Indian gene pool line Varuna and was found to confer complete resistance to all the isolates. This is the first white rust resistance-conferring gene described from *Brassica* species and has been named *BjuWRR1*. Allelic variants of the gene in *B. juncea* germplasm and orthologues in the Brassicaceae genomes were studied to understand the evolutionary dynamics of the *BjuWRR1* gene.

**Highlight:** *BjuWRR1*, a CNL type R gene, was identified from an east European gene pool line of *Brassica juncea* and validated for conferring resistance to white rust by genetic transformation.

## Introduction

White rust, caused by an oomycete pathogen *Albugo candida* (Pers.) Kuntze is a serious disease of economically important *Brassica* species. *A. candida* is an obligate biotrophic parasite and rated amongst the top ten oomycete pathogens based on its scientific and economic importance (Kamoun *et al.*, 2015). White rust has been reported to cause significant yield losses in *B. juncea* – anywhere from 20% to 60% in India, Australia and Canada (Rimmer *et al.*, 2000; Kaur *et al.*, 2008; Awasthi *et al.*, 2012). The impact of the disease is highest in the Indian subcontinent as mustard is grown in more than five million hectares of land. Almost all the released lines grown commercially in India are susceptible to the disease (Li *et al.*, 2008; Panjabi-Massand *et al.*, 2010; Awasthi *et al.*, 2012) The incidence of white rust is more pronounced after the rains – as a consequence, yield gains occurring from additional moisture under the dryland agronomic conditions are lost due to high incidence of the disease.

White rust disease is characterized by the formation of white to cream-colored zoosporangial pustules on all the aerial parts of the plants such as cotyledons, leaves, stems and inflorescences (Saharan *et al.*, 2014). Systemic spread of the pathogen from an early infection of the vegetative parts to the inflorescence causes hypertrophy of the floral tissues resulting in the formation of staghead galls that produce little or no seed (Verma and Petrie, 1980; Meena *et al.*, 2014). The disease can be controlled with fungicides (Meena and Jain 2002); however, the use of chemicals, besides being ecologically undesirable increases the input costs in mustard cultivation adversely affecting incomes of the small and marginal farmers. Exploitation of the host-plant resistance would be the most efficient, environment-friendly and cost-effective method of controlling white rust disease.

Several efforts have been made for the identification of loci in *Brassica* species conferring resistance to different races of *A. candida*. Such loci have been mapped in B. *rapa* (Kole *et al.*, 1996, 2002), B. *napus* (Ferreira *et al.*, 1995) and B. *juncea* (Cheung *et al.*, 1998; Mukherjee *et al.*, 2001; Panjabi-Massand *et al.*, 2010). However, none of the mapping studies in Brassicas have been taken forward to characterize the genes involved with resistance to white rust. To date, only two R genes, *RAC1* and *WRR4* conferring resistance to *A. candida* have been identified and cloned from the model crucifer species *Arabidopsis thaliana* (Borhan *et al.*, 2004, 2008). Both these genes belong to the TIR-NB-LRR subfamily. *WRR4* gene was introduced into *B. juncea* line Cutlass, an east European gene pool line susceptible to *A. candida* isolate used in the study, by genetic transformation and was found to confer significant levels of resistance (Borhan *et al.*, 2010).

In a previous study, two independent loci conferring resistance to race 2 of *A. candida* (isolate AcB1) were mapped in two east European *B. juncea* gene pool lines, Heera and Donskaja-IV using two F_1_ DH mapping populations – VH (Varuna x Heera) and TD (TM-4 x Donskaja-IV), Varuna and TM-4 being the susceptible Indian gene pool lines (Panjabi-Massand *et al.*, 2010). In Heera, a partial resistance-conferring locus AcB1-A4.1 was mapped to the linkage group A04, while in Donskaja-IV, a complete resistance-conferring locus AcB1-A5.1 was mapped to the linkage group A05.

In the present study, locus AcB1-A5.1 mapped in Donskaja-IV has been studied further to characterize the resistance-conferring gene present in the mapping interval. A new differential set of *Brassica* species has been developed and tested with six isolates of *A. candida* isolated from different mustard growing regions of north India. A combination of positional cloning and candidate gene approach was used to identify a white rust resistance-conferring R gene named *BjuA5.WRR.a1* (in short *BjuWRR1*) that was validated by genetic transformation of a susceptible line. We further report on the structure of the *BjuWRR1* gene, its allelic variants in *B. juncea* germplasm and its orthologues in the sequenced genomes of different species belonging to the family Brassicaceae.

## Materials and methods

### Plant materials

Germplasm used in the study is listed in Table 1 (more details are provided in Supplementary Table S1). All the *B. juncea* lines used and experimental lines developed in the study were maintained by strict selfing. Self-incompatible lines of *B. rapa* and *B. nigra* were maintained by sib-mating.

**Table 1.** Disease reactions of *Brassica juncea* lines and *Brassica* species, AcB1-A5.1 NILs and Varuna transgenic lines containing the *BjuWRR1* gene to *Albugo candida* isolates collected from six different locations of north India.

|  |  | Isolates/ Disease scoring* |  |  |  |  |  |
| --- | --- | --- | --- | --- | --- | --- | --- |
|  |  | Pantnagar<br>(AcP1) | Meerut<br>(AcM1) | Samastipur<br>(AcS1) | Hisar<br>(AcH1) | Bharatpur<br>(AcB1) | Alwar<br>(AcA1) |
| **Reference lines** |  |  |  |  |  |  |  |
| *B. juncea* | Varuna | S | S | S | S | S | S |
|  | Heera | S | S | S | R | R | S |
|  | Donskaja-IV | R | R | R | R | R | R |
|  | Tumida | R | MS | R | R | R | R |
|  | Cutlass | R | R | R | R | R | R |
|  | Skorospieka | S | S | S | S | S | S |
| *B. rapa* | YSPB | R | MS | R | R | R | R |
|  | Chiifu-401 | S | R | S | S | MR | R |
|  | Candle | MR | MR | MR | MR | R | MR |
|  | Pant toria | S | S | S | S | S | S |
| *B. nigra* | Sangam | S | S | S | S | S | S |
|  | BN2782 | MR | R | MR | MR | R | MR |
| **Near isogenic lines** |  |  |  |  |  |  |  |
| Varuna | Varuna-936-266 | R | R | R | R | R | R |
|  | Varuna-865-245 | R | R | R | R | R | R |
|  | Varuna-583-37 | R | R | R | R | R | R |
| Pusa bold | PB-A5-223 | R | R | R | R | R | R |
| Pusa jai kisan | PJK-A5-63 | R | R | R | R | R | R |
| Rohini | Rohini-A5-467 | R | R | R | R | R | R |
| **Transgenic lines** |  |  |  |  |  |  |  |
| (Varuna) | BjuWRR1-2 | R | R | R | R | R | R |
|  | BjuWRR1-3 | R | R | R | R | R | R |

\*Disease scoring is represented by four classes: R, resistant (scores 0 and 1); MR, moderately resistant (score 3); MS, moderately susceptible (score 5) and S, susceptible (scores 7 and 9).

### Development of AcB1-A5.1 near-isogenic lines (NILs)

NILs for white rust resistance-conferring locus AcB1-A5.1 were developed in susceptible Indian lines Varuna, Pusa bold, Pusa jai kisan and Rohini by marker-assisted-backcross breeding (MAB). Locus AcB1-A5.1 was introgressed from the donor parent, Donskaja-IV into susceptible lines through positive selection by a PCR-based Intron polymorphism (IP) marker At2g36360 (Panjabi-Massand *et al.*, 2010) tightly linked to the resistance phenotype and negative selection by IP marker At2g44970 on one side and SSR marker At2g34700 on the other side. Detailed information on the markers is available in our previous studies (Panjabi *et al.*, 2008; Dhaka *et al.*, 2017; Rout *et al.*, 2018). Nucleotide sequences of the primers used in the present study for the markers are given in Supplementary Table S2. The entire scheme took five generations of back-crossing followed by selfing to develop homozygous NILs.

### *A. candida* isolates, infection and disease evaluation

White rust infected *B. juncea* leaves were collected from five different mustard growing regions of north India (Pantnagar, Meerut, Samastipur, Hisar and Alwar) and were purified by performing five passages of inoculations for each isolate on a susceptible *B. juncea* line, Varuna. In each passage, zoosporangia collected from a single pustule were suspended in 200 μL of sterilized double distilled water followed by drop inoculation on cotyledons of seven-day-old seedlings of Varuna. Inoculations were performed in a contained white rust infection chamber maintained at a relative humidity of around 90% and 18°C of temperature with a 16-h light/8-h dark cycle. After repeated passages of single pustules, the isolates were maintained on Varuna and the purified *A. candida* isolates were named as AcP1 (Pantnagar), AcM1 (Meerut), AcS1 (Samastipur), AcH1 (Hisar) and AcA1 (Alwar).

The above-mentioned isolates along with a previously reported isolate from Bharatpur, AcB1 (Panjabi-Massand *et al.*, 2010) were evaluated for infectivity on some core germplasm which will be referred to as the differential set. Seeds were germinated in pots containing sterilized soil and soilrite in growth chambers (Conviron, Canada) maintained at 20°C with a 10-h light/14-h dark cycle and 70% relative humidity. Inoculations on cotyledons of seven-day-old seedlings were performed following the procedure described by Panjabi-Massand *et al*. (2010). For the evaluation of white rust response; disease scoring was carried out following the procedures described by Williams (1985) and Panjabi-Massand *et al*. (2010). Disease scoring was done fourteen days post-inoculation and the disease phenotypes were rated as 0, 1, 3, 5, 7 and 9 as described by Panjabi-Massand *et al*. (2010) and further categorized into four classes, scores 0 and 1 – resistant (R), score 3 – moderately resistant (MR), score 5 – moderately susceptible (MS) and scores 7 and 9 – susceptible (S). Three independent infection assays were carried out and every experiment consisted of at least five seedlings of each of the tested lines.

### BAC isolation, sequencing and annotation

A 5x pooled and amplified HindIII BAC library was developed (Bio S&T, Canada) using genomic DNA extracted from young leaves of Donskaja-IV. Partially-digested, size-selected DNA fragments (150-350kb) were ligated into HindIII digested vector pIndigoBAC-5 (Epicentre, USA) and transformed into *E. coli* DH10B cells (Invitrogen, USA). Colonies were distributed into 1 x 96 well plate (Pooled BAC library). Each well contained about 573 primary clones (Total clones > 55,000). The BAC library was screened using PCR based IP marker At2g36360. Two additional IP markers (At2g36480 and At2g36310) flanking At2g36360 were developed using sequence information of *B. rapa* line Chiifu-401 obtained from the BRAD database (Cheng *et al.*, 2011). BAC clones giving amplifications with all the three markers were sequenced on a PacBio RSII instrument based on single-molecule realtime (SMRT) technology followed by assembly of the reads using a hierarchical genome assembly process (Amplicon Express, USA).

BAC sequences were annotated for ORFs using Augustus (Stanke *et al.*, 2004), a web server for gene prediction in eukaryotes. ORFs were screened for conserved domains using NCBI Batch CD-Search (Marchler-Bauer and Bryant, 2004; Marchler-Bauer *et al.*, 2011). The domains of BjuWRR1 protein were also predicted using an online tool – InterPro (Finn *et al.*, 2017) and COILS (ExPASy Bioinformatics Resource portal). The conserved motifs of the NB domain, described by Meyers *et al*. (2003), were identified manually. Molecular weight and isoelectric point of the protein was computed by Compute pI/MW online (ExPASy Bioinformatics Resource portal, https://web.expasy.org/compute_pi/).

### Vector construction and genetic transformation of *B. juncea*

PCR amplification of the candidate gene using BAC clone as a template was carried out with Q5 High-Fidelity DNA Polymerase (New England Biolabs, USA). The amplified product was cloned into a modified binary vector pCGMCP22 (Arumugam *et al.*, 2007) containing *bar* gene conferring resistance to phosphinothricin (PPT) as a selectable marker between the *loxP* sites. The *bar* gene was driven by the CaMV35S promoter with an AMV leader sequence (Jobling and Gehrke, 1987) and ocspA as the terminator. The modified vector was digested with restriction enzymes HindIII and PstI (New England Biolabs, USA) and amplified product was cloned using In-Fusion HD Cloning Kit (Clontech, USA) followed by sequencing on a 3730 DNA Analyzer (Applied Biosystems, USA).

The binary vector containing the cloned gene was transferred into disarmed *Agrobacterium tumefaciens* strain GV3101 by electroporation. Genetic transformation of *B. juncea* line Varuna was carried out using hypocotyl explants following a protocol described by Mehra *et al*. (2000). Positive transformants were selected *in vitro* on a medium containing PPT at a concentration of 10 mg L^-1^. For segregation analysis of the progeny of the T_0_ transgenics for resistance and susceptibility to PPT, a concentration of 200 mg L^-1^ was used in the field.

For Southern blot analysis, genomic DNA was isolated from well-expanded leaves collected from the field grown transgenic lines (T_0_) following the protocol of Rogers and Bendich (1994). 6 μg of DNA was digested with restriction enzyme EcoRI (New England Biolabs, USA). Digested DNA was separated on 0.8% agarose gel and blotted onto Hybond-N+ nylon membrane (Amersham Pharmacia Biotech, USA). The blots were hybridized with probes that were labeled using PCR DIG Probe Synthesis Kit and hybridization was detected with DIG Luminescent Detection Kit following the manufacturer’s instructions (Roche, Switzerland).

### Gene expression and cDNA analysis

For studying changes in transcript accumulation in response to pathogen infection, cotyledons of one-week-old seedlings of Donskaja-IV were inoculated with 10 μL of sporangial suspension (2 × 10^4^ sporangia mL^-1^) of *A. candida* isolate AcB1 using drop inoculation method. The cotyledon samples inoculated with the pathogen and distilled sterile water (mock) were collected 4h, 48h, 72h and 14 days after the inoculations – with four seedlings for each sample. Total RNA was extracted from the collected samples using Spectrum Plant Total RNA Kit (Sigma-Aldrich, USA) and was treated with RNase-free DNaseI (Qiagen, Netherlands) to remove any contaminating genomic DNA. 2 μg of total RNA was used to synthesize the first-strand cDNA using High-Capacity cDNA Reverse Transcription Kit (Applied Biosystems, USA) according to the manufacturer’s instructions. Real-time quantitative reverse transcription-PCR (qRT-PCR) analyses were carried out using a PowerUp SYBR Green Master Mix (Applied Biosystems, USA) on QuantStudio 6 Flex Real-Time PCR System (Applied Biosystems, USA). Comparisons of relative gene expression were made between the pathogen and mock-inoculated samples collected at the same time points. The ubiquitin gene, UBQ9 (Chandna *et al.*, 2012) was used as an internal control. The 2^-ΔΔ*C*^_T_ method (Livak and Schmittgen, 2001) was followed in evaluating the relative gene expression levels.

To obtain cDNA sequence, RT-PCR was carried out using total RNA isolated from one-week-old cotyledons of Donskaja-IV. The first strand cDNA was synthesized as described above and was used as a template for PCR reaction using Phusion High-Fidelity DNA Polymerase (Thermo Fisher Scientific, USA) with a gene-specific primer pair. The PCR product was cloned in the pGEM-T-Easy vector (Promega, USA) and sequenced on a 3730 DNA Analyzer (Applied Biosystems, USA).

### Sequence information of syntenic regions in *B. juncea* lines and sequenced Brassicaceae genomes

Sequence of the syntenous region in line Heera was obtained by screening a genomic DNA BAC library of the line reported earlier (Sharma *et al.*, 2014). Identified BAC clone was sequenced by the procedure reported for the Donskaja-IV BAC. Sequence of line Varuna was obtained from a SMRT based genomic assembly (unpublished data). Sequences present in Cutlass, Skorospieka, Kranti, *B. nigra* (Sangam) and the B genome of *B. juncea* (Donskaja-IV) were obtained from Illumina short-read sequencing of the genomes of these lines (unpublished data); sequence of Tumida was downloaded from the BRAD database (Wang *et al.*, 2015). Sequence information of other members of the Brassicaceae family were obtained from the BRAD database. Multiple sequence alignment of the alleles was done with ClustalW using MegAlign software (DNASTAR, USA).

## Results

### Testing of *A. candida* isolates on a differential set

*A. candida* isolates – AcP1, AcM1, AcS1, AcH1, AcB1 and AcA1 collected from different regions of north India were evaluated for their infectivity on a set of differentials consisting of six lines of *B. juncea* (AABB), four of *B. rapa* (AA) and two of *B. nigra* (BB) (Table 1). The NILs, containing the locus AcB1-A5.1, developed in four susceptible Indian *B. juncea* lines – Varuna, Pusa bold, Pusa jai kisan and Rohini were also tested for their response against these isolates. Disease response was recorded in four classes: R (resistant), MR (moderately resistant), MS (moderately susceptible) and S (susceptible) as described in the materials and methods section. The infection results showed variations in the disease response to the collected isolates on the tested lines except for the isolates AcP1 and AcS1 which showed similar disease response. This analysis suggested that apart from the isolates AcP1 and AcS1, all the other collected isolates differ from each other in their effector sequences and/or effector profile. Donskaja-IV and Cutlass were the only lines completely resistant to all the tested isolates and would, therefore, serve as vital sources for resistance gene(s) that can be deployed for complete resistance. The AcB1-A5.1 NILs in all the varietal backgrounds that are otherwise highly susceptible, showed complete resistance – similar to that present in the donor parent Donskaja-IV making this locus interesting for further genetic dissection and molecular characterization.

### Identification of candidate gene(s) in AcB1-A5.1 locus of Donskaja-IV

The white rust resistance-conferring locus, AcB1-A5.1, identified in Donskaja-IV was characterized by screening Donskaja-IV BAC library using PCR-based marker – At2g36360 tightly linked to the resistant trait and the flanking markers – At2g36480 and At2g36310. Two BAC clones 7B and 12E of sizes 152.1 kb and 186.1 kb, respectively, were identified and sequenced. As the sequence information of both the BACs was similar, only the clone 7Bwas characterized further. Twenty-eight protein encoding sequences were predicted in the genomic sequence of the identified BAC (Fig. 1). An analysis of the conserved protein domains in the twenty-eight ORFs (Supplementary Table S3), revealed the presence of eight ORFs (ORF1, ORF4, ORF7, 0RF20, ORF22, ORF23, ORF24 and ORF26) encoding for Homeobox, CBS, WRC, QLQ, CC-NB-LRR, Kelch, protein kinase, and A20/AN1-like Zinc finger domain-containing proteins that could be involved in conferring resistance to pathogens (Coego *et al.*, 2005; Wang *et al.*, 2005; Liu *et al.*, 2008; Hewezi *et al.*, 2012; Liu *et al.*, 2014 b; Tyagi *et al.*, 2014; Zuo *et al.*, 2015; Ma *et al.*, 2015; Mou *et al.*, 2015; Omidbakhshfard *et al.*, 2015; Wang *et al.*, 2017). Since a majority of the cloned R genes till date belong to the NB-LRR class (Kourelis and van der Hoorn, 2018), among the eight potential genes for conferring resistance, ORF22 encoding a CC-NB-LRR domain-containing protein as predicted by the online tool Interpro was considered to be the most likely candidate for conferring resistance to white rust in Donskaja-IV. We named the CC-NB-LRR encoding gene – *BjuA5.WRR.a1* following the standardized gene nomenclature suggested for the *Brassica* genus by Ostergaard and King (2008). However, we will hereafter refer to the gene as *BjuWRR1* – as it is the first gene to be cloned from *B. juncea* that confers resistance to white rust.

**Fig. 1.**
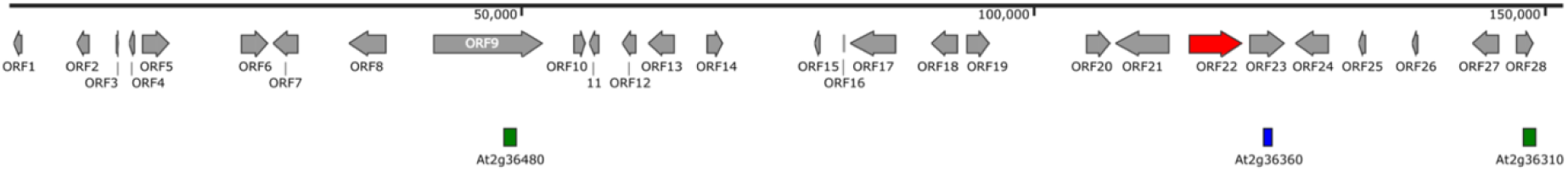
Physical map of the BAC clone spanning the AcB1-A5.1 region of Donskaja-IV. The BAC clone (152.1 kb) was identified by screening Donskaja-IV BAC library using a tightly linked IP marker At2g36360 (indicated by a blue box) and flanking markers At2g36480 and At2g36310 (indicated by green boxes). Open reading frames (ORFs) predicted in the region are shown in grey with forward or reverse orientation of the genes being depicted by the arrowheads. Red arrow (ORF22) is the predicted CC–NB–LRR protein encoding candidate R gene, *BjuWRR1* present next to the tightly linked marker.

### Validation of *BjuWRR1* as white rust resistance-conferring gene by genetic transformation

To validate the role of the candidate gene *BjuWRR1* in conferring white rust resistance, genetic transformation experiments were carried out. *BjuWRR1* was amplified from the BAC clone 7B. PCR primers were designed (BjuWRR1inf, Supplementary Table S2) from the regions 1.5 kb upstream of the start codon and 0.9 kb downstream of the stop codon of *BjuWRR1* so as to encompass the gene’s native promoter and terminator sequences. Using BAC DNA as the template, a 7.7 kb genomic DNA fragment was amplified and cloned in the pCGMCP22-bar(lox) binary vector. The vector (Supplementary Fig. S1) was used to transform a highly susceptible line of *B. juncea*, Varuna (Table 1) belonging to the Indian gene pool of *B. juncea*. Twenty-eight independent transgenic lines were screened for single copy insertions of the *bar* and *BjuWRR1* genes by performing Southern hybridizations with the probes developed using primer pairs DIG-bar and DIG-BjuWRR1 (Supplementary Table S2). The position from where the probes were designed for Southern hybridization and EcoR1 sites within the T-DNA are shown in the Supplementary Fig. S2. The number of copies inserted in the transgenic lines varied (Supplementary Table S4). Transgenic lines BjuWRR1-2 and BjuWRR1-3 that contained the single copy insertion of the T-DNA containing both the *bar* and *BjuWRR1* gene cassettes, were used for further analysis. These two lines also showed a 3:1 segregation ratio for resistance and susceptibility to PPT in the progeny of the T0 plants (Supplementary Table S5).

Infection assays were carried out using *A. candida* isolate AcB1(earlier used for mapping the locus) on the T_1_ segregants of BjuWRR1-2 and BjuWRR1-3 transgenic lines. The untransformed Varuna line which served as the control was highly susceptible, whereas the T_1_ of the two transgenic lines segregated for complete resistance (as in Donskaja-IV) or susceptibility (as in Varuna). Segregation for resistance and susceptibility to white rust disease in T_1_ generation was confirmed by performing PCR, using *bar* specific (Bar, Supplementary Table S2) and *BjuWRR1* specific (BjuWRR1, Supplementary Table S2) primer pairs. Resistance in T_1_ lines to isolate AcB1co-segregated with the presence of the *bar* and *BjuWRR1* genes (Fig. 2). Complete resistance to isolate AcB1 in the Varuna transgenics containing the *BjuWRR1* gene validated the role of this CNL type R gene in conferring complete resistance to the white rust disease.

**Fig. 2.**
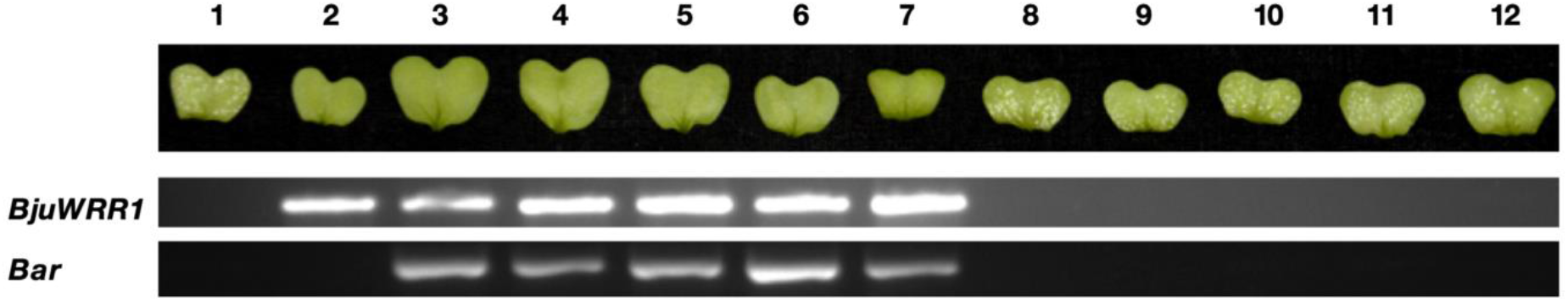
Infection assays carried out on the susceptible *B. juncea* line Varuna transformed with the *BjuWRR1* gene of Donskaja-IV with *A. candida* isolate AcB1 (top) and molecular analysis using primers specific to the *BjuWRR1* and *bar* genes (bottom). Numbers 1 and 2 represent *B. juncea* lines Varuna and Donskaja-IV, respectively; numbers from 3 to 7 and 8 to 12 represent resistant and susceptible T1 progenies, respectively, of the transgenic line BjuWRR1-2.

### Response of transgenic lines to multiple *A. candida* isolates

Infection experiments on Donskaja-IV and NILs for locus AcB1-A5.1 had revealed that all the susceptible lines containing locus AcB1-A5.1 were completely resistant to a diverse set of *A. candida* isolates (Table 1). We checked the extent of white rust resistance achieved by the *BjuWRR1* gene alone in the transgenic events BjuWRR1-2 and BjuWRR1-3 to all the isolates. Infection assays were performed on T_1_ segregants of the transgenic lines. Complete resistance co-segregated with the presence of the *BjuWRR1* transgene. The results showed that *BjuWRR1* conferred complete resistance against all the isolates in both the transgenic lines as was the case in the donor parent, Donskaja-IV (Table 1). This confirmed *BjuWRR1* as the key gene in the AcB1-A5.1 locus that is involved with conferring white rust resistance.

### Characterization of *Bju WRR1*

The cDNA of *BjuWRR1* was cloned from the resistant *B. juncea* variety Donskaja-IV using gene-specific primer pair (BjuWRR1seq, Supplementary Table S2). A comparison of the genomic and cDNA sequences of *BjuWRR1* gene showed a 2739 bp coding region (Fig. 3A) that encoded a polypeptide of 912 amino acid residues with an estimated molecular weight of 105.6 kDa and a theoretical isoelectric point (pI) of 7.8. Analysis of the protein sequences showed that *BjuWRR1* encoded a protein that contained canonical CC, NB and LRR domains. Further, several conserved motifs present in the NB domain reported by Meyers *et al*. (2003) that include P-loop (residues 183 to 201), RNBS-A (residues 206 to 234), Kinase 2 (residues 261 to 271), RNBS-B (residues 287 to 301), RNBS-C (residues 312 to 329), GLPL (residues 350 to 365), RNBS-D (residues 418 to 446) and MHDV (residues 486 to 500) could be identified by manual inspection (Fig. 3B). Thus, the R gene *BjuWRR1* encoded a typical CC-NB-LRR protein.

**Fig. 3.**
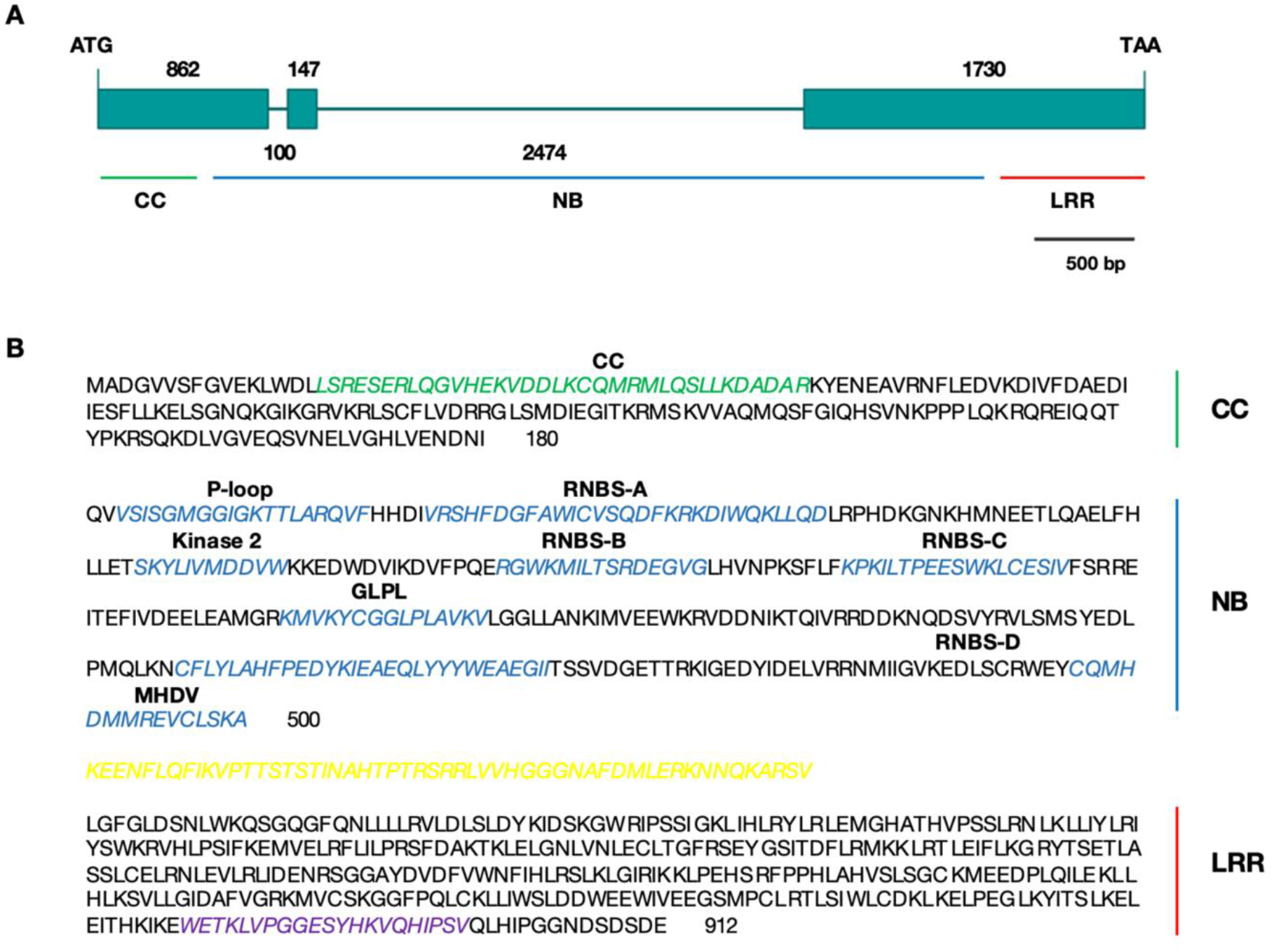
Structure and amino acid sequence of BjuWRR1. (A) Gene structure of *BjuWRR1* – ATG and TAA indicate the translation start and stop codons, respectively. The green boxes represent exons and lines denote introns with length described by the number of base pairs. (B) Deduced protein sequence of the *BjuWRR1* gene. The coiled-coil (CC) and leucine-rich repeat (LRR) domains are shown in green and red, respectively. The conserved motifs (P-loop, RNBS-A, Kinase 2, RNBS-B, RNBS-C, GLPL, RNBS-D and MHDV) of the nucleotide binding (NB) region are shown in blue. The NB-LRR linker and the conserved C-terminus sequences are highlighted in yellow and purple, respectively.

### Expression analysis of *Bju WRR1*

To determine whether expression of *BjuWRR1* in Donskaja-IV varied in response to a challenge with the pathogen, we evaluated the expression of *BjuWRR1* at different time points post-inoculation with *A. candida* isolate AcB1 using qRT-PCR. A primer pair (RT-BjuWRR1, Supplementary Table S2) specific to *BjuWRR1* was designed from the intron-spanning exons (positions of the primers are shown in Fig. 4A). The *BjuWRR1* expression was detected both before and after inoculation, indicating that this gene is constitutively expressed. The relative expression of the gene in the infected seedlings with the mock-inoculated seedlings of Donskaja-IV at different time points showed no significant change in the expression of the gene indicating that the levels of the mRNA that were present preinfection were sufficient for providing resistance to the white rust disease (Fig. 4B).

**Fig. 4.**
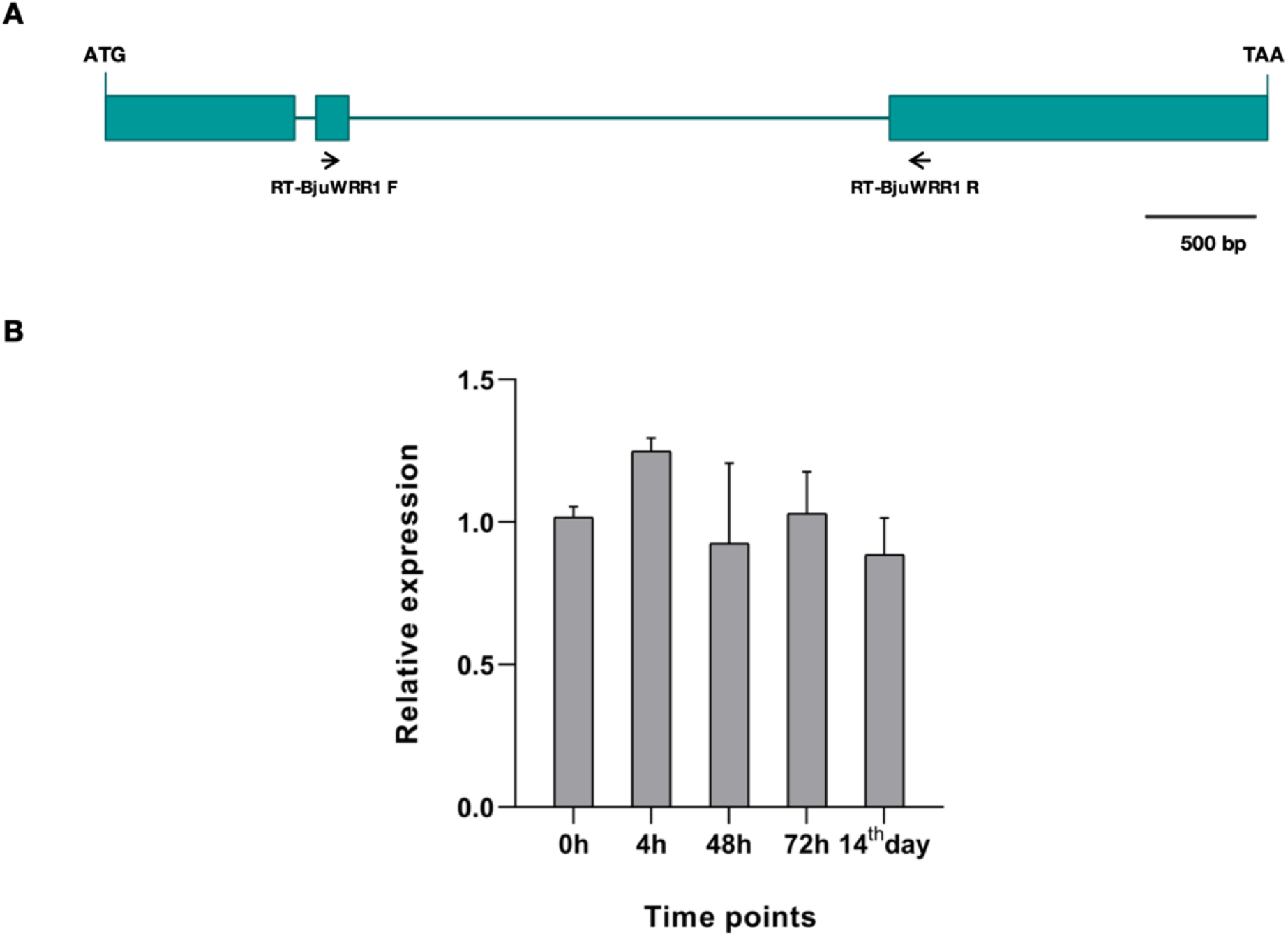
Schematic representation of the placement of primers used in qRT-PCR for expression analysis of *BjuWRR1* in Donskaja-IV – pre and post-inoculation with *Albugo candida* isolate AcB1. (A) Representation of the *BjuWRR1* gene highlighting the position from where the primer pair RT-BjuWRR1 was designed. (B) Expression analysis of *BjuWRR1* in seven-day-old seedlings of Donskaja-IV over different time intervals after inoculation with *Albugo candida* isolate AcB1. Values represent the average of three independent biological replicates. Bars represent standard errors.

### Allelic variants of gene *Bju WRR1* in *B. juncea* lines

Allelic diversity in the *BjuWRR1* gene was studied in six lines of *B. juncea*, two Indian gene pool lines that have been extensively grown in India (Varuna and Kranti), three from the east European gene pool (Heera, Cutlass and Skorospieka) and one vegetable type from China (Tumida). The structural organization of different alleles, from the start to stop codon, has been shown in Fig. 5. Expression of *BjuWRR1* alleles in Varuna, Heera and Tumida each belonging to a different gene pool were examined by performing RT-PCR using three different primer pairs (CDSE1-2, CDSE2-3, CDSE3-4, Supplementary Table S2) designed from the predicted exonic regions spanning the intron (position of the primers used is shown in Fig. 5). The results validated the *in silico* prediction of the exons and constitutive expression of the alleles. Based on the comparison of coding sequences and the encoded amino acids (Supplementary Fig. S3), three types of alleles were found in the analyzed lines. Donskaja-IV_*BjuWRR1* (allelic variant 1) and Cutlass_*BjuWRR1* (allelic variant 2) were more similar compared to the third allelic variant that was conserved in the remaining lines. Donskaja-IV_*BjuWRR1* and Cutlass_*BjuWRR1* showed an amino acid identity of 99%. Cutlass_*BjuWRR1* protein would be shorter due to a premature stop codon in the allele leading to a loss of 115 amino acids at the C-terminal end compared to the amino acid sequence of Donskaja-IV_BjuWRR1. Surprisingly, the third type of allele, V/H/S/K/T*_BjuWRR1* present in Varuna, Heera, Skorospieka, Kranti and Tumida showed identical coding and amino acid sequences with minor variations in the sequence of the intronic regions. V/H/S/K/T *BjuWRR1* showed the nucleotide and amino acid identity of 92% and 86% respectively with Donskaja-IV_*BjuWRR1*. A comparison of the synonymous and non-synonymous substitutions in different alleles, with Donskaja-IV_BjuWRR1 as the reference sequence, showed that the non-synonymous substitutions between Donskaja-IV_*BjuWRR1* and V/H/S/K/T_*BjuWRR1* outnumber the synonymous substitutions (Fig. 6).

**Fig. 5.**
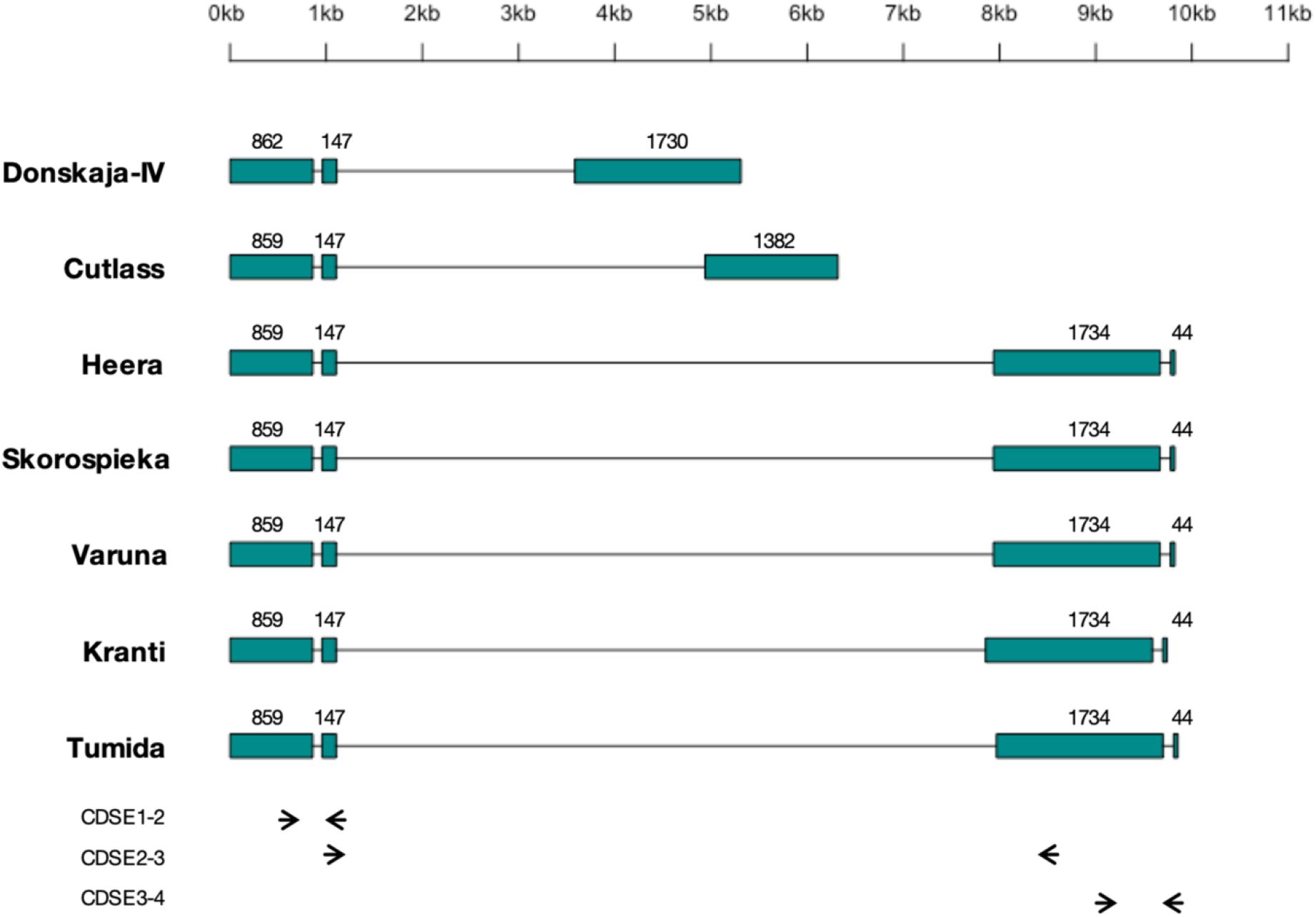
Structure of allelic variants of *BjuWRR1* gene in different lines of *B. juncea*. The boxes represent exons and lines indicate introns with length denoted in base pairs. All the alleles have been drawn from the start codon to the stop codon. Predicted exons of *BjuWRR1* alleles in Varuna, Heera and Tumida were validated by performing PCR using respective cDNAs as template and three pairs of intron spanning exon specific primers (CDSE1-2, CDSE2-3 and CDSE3-4) depicted by arrows.

**Fig. 6.**
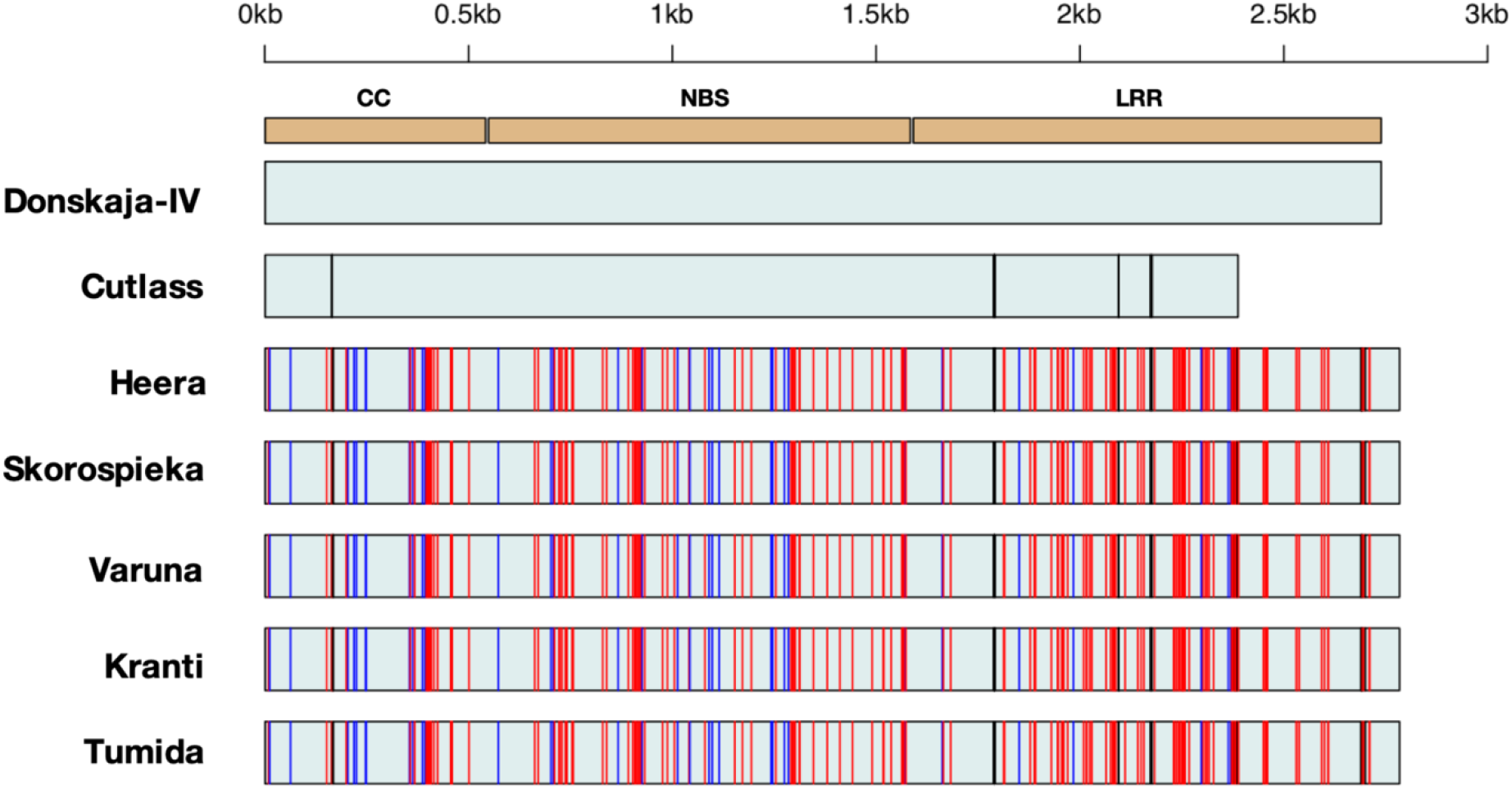
Comparison of the nucleotide sequences of the coding regions of *BjuWRR1* alleles present in different *B. juncea* lines. The sequence of Donskaja-IV_B;uWRR1 was used as a reference with which other sequences were compared. Synonymous and Non-synonymous substitutions are highlighted in blue and red vertical lines, respectively, whereas black lines denote deletions.

### Evolution of *BjuWRR1* across species

*BjuWRR1* is present in the least fractionated (LF) J-block on linkage group A05 of Donskaja-IV. We scanned the syntenic regions of the sequenced species of Brassicaceae family – *B. rapa, B. nigra, B. oleracea, B. napus, B. juncea, A. thaliana, A. arabicum, A. lyrata, Capsella rubella, C. sativa, Sisymbrium irio, Thellungiella halophile* and *T. salsuginea*. For finding *BjuWRR1* orthologues in different species we focused on the syntenic regions containing the genes encoding for 1, 4-alpha-glucan branching enzyme (At2g36390), kelch repeat-containing protein (At2g36360) and a protein kinase (At2g36350) that flank the *BjuWRR1* gene in Donskaja-IV. Details of the species used in the analysis, source of the sequences, description of the flanking genes and *BjuWRR1* orthologues have been described in the Supplementary Table S6. Three major haplotypes were observed at the *BjuWRR1* locus based on the presence/absence and copy number variation of the CNL type genes at this locus (Fig. 7). An absence of a CNL type gene was classified as haplotype H1 and was characteristic of *A. thaliana*, *A. arabicum*, *A. lyrata*, *C. rubella*, *C. sativa*, *S. irio*, *T. halophile* and *T. salsuginea*. Presence of a simple genetic locus with either complete, incomplete or remnants of a single CNL gene was classified as haplotype H2 which was characteristic of *B. nigra, B. oleracea*, A and B subgenomes of *B. juncea* and C subgenome of *B. napus*. Presence of a complex locus with more than one CNL gene was classified as haplotype H3 and was observed in *B. rapa* and the A subgenome of *B. napus*. Comparison of the *BjuWRR1* syntenic region in the triplicated ‘A’ subgenomes (LF, MF1 and MF2) of Donskaja-IV suggested that *BjuWRR1* gene is specific to LF and is not present, even as a remnant, in the MF1 and MF2 subgenomes.

**Fig. 7.**
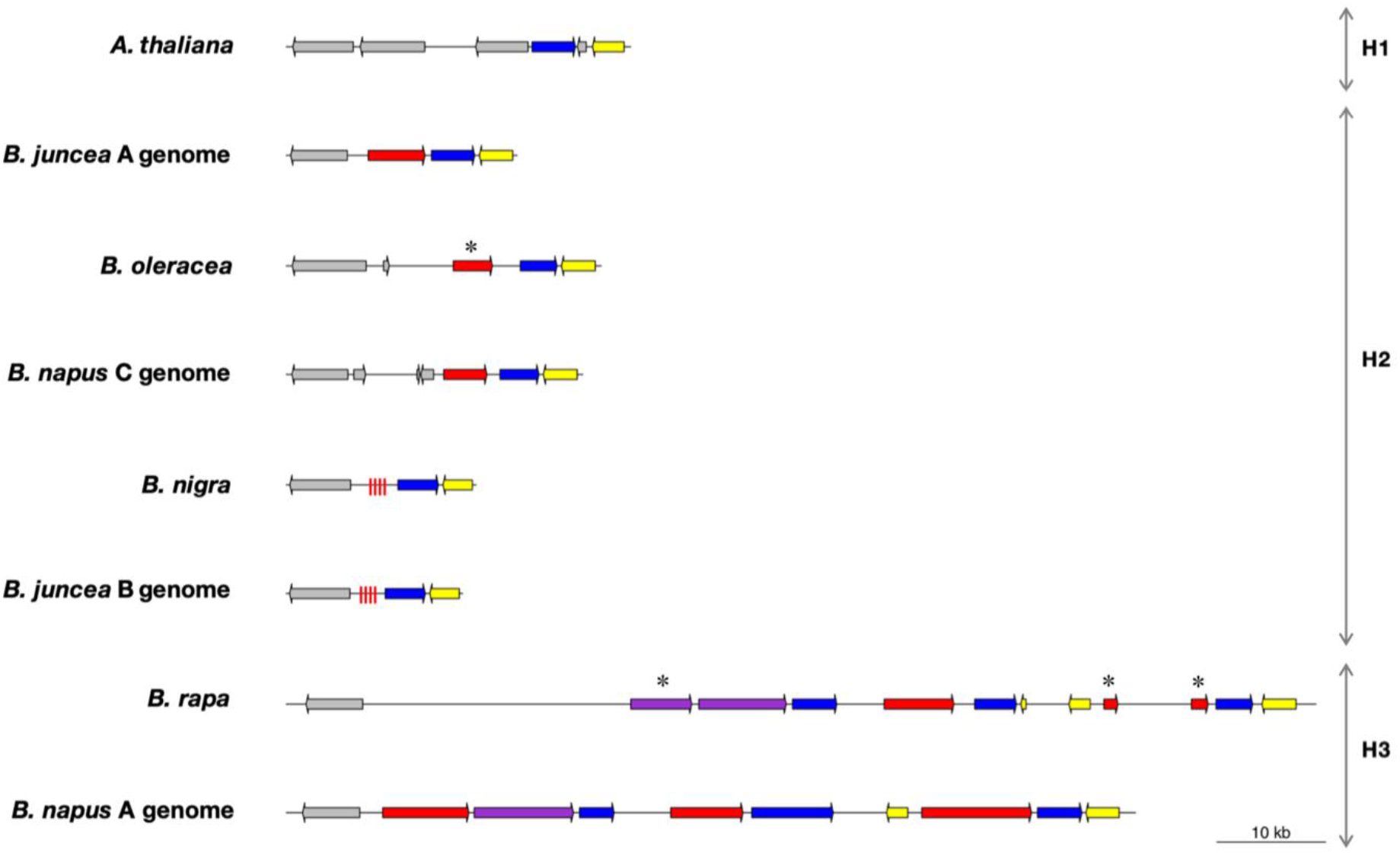
Organization of *BjuWRR1* gene orthologues in some of the sequenced members of the Brassicaceae family. The three haplotypes (H1, H2 and H3) observed in this locus are shown on the right. CNL genes are represented in red. Kelch repeat and Ser/thr-protein kinase encoding genes are highlighted in blue and yellow respectively. Genes represented in purple are predicted to encode for proteins containing both CNL domains and kelch repeat. Red dashed lines indicate remnants of a CNL type gene. Asterisks highlight the predicted CNL pseudogenes.

To gain a better insight into the *BjuWRR1* orthologues in the AcB1-A5.1 region of the species with H2 and H3 haplotypes, all the predicted R-genes in the syntenic regions were manually inspected and classified as full-length ORFs encoding CNL proteins or pseudogenes. Of the 12 orthologues that were identified, four were truncated either at 5’ or at 3’ termini leading to absence of the CC, NB and/or LRR domains. In the absence of functional data for these genes, it cannot be inferred with certainty whether these are pseudogenes or are functional genes.

### Development of allele-specific marker for Donskaja-IV_*BjuWRR1*

The genomic sequence of Donskaja-IV_*BjuWRR1* showed that exon 2 shared significant similarity with the corresponding region of Varuna_*BjuWRR1*, but not much similarity was found in the intron 2 region. A common forward primer, F-DV from exon 2 and specific reverse primers R-D for Donskaja-IV and R-V for Varuna from the intron 2 region were designed (Fig. 8A). Using the three primers, a 761-bp fragment was amplified from Varuna while a 366-bp fragment was amplified from Donskaja-IV. These primers were tested on different Indian *B. juncea* lines namely – TM-4, Kranti, Pusa bold, Pusa jai kisan, Rohini, RH30 and Rajat and gave an amplification similar to that in Varuna.

**Fig. 8.**
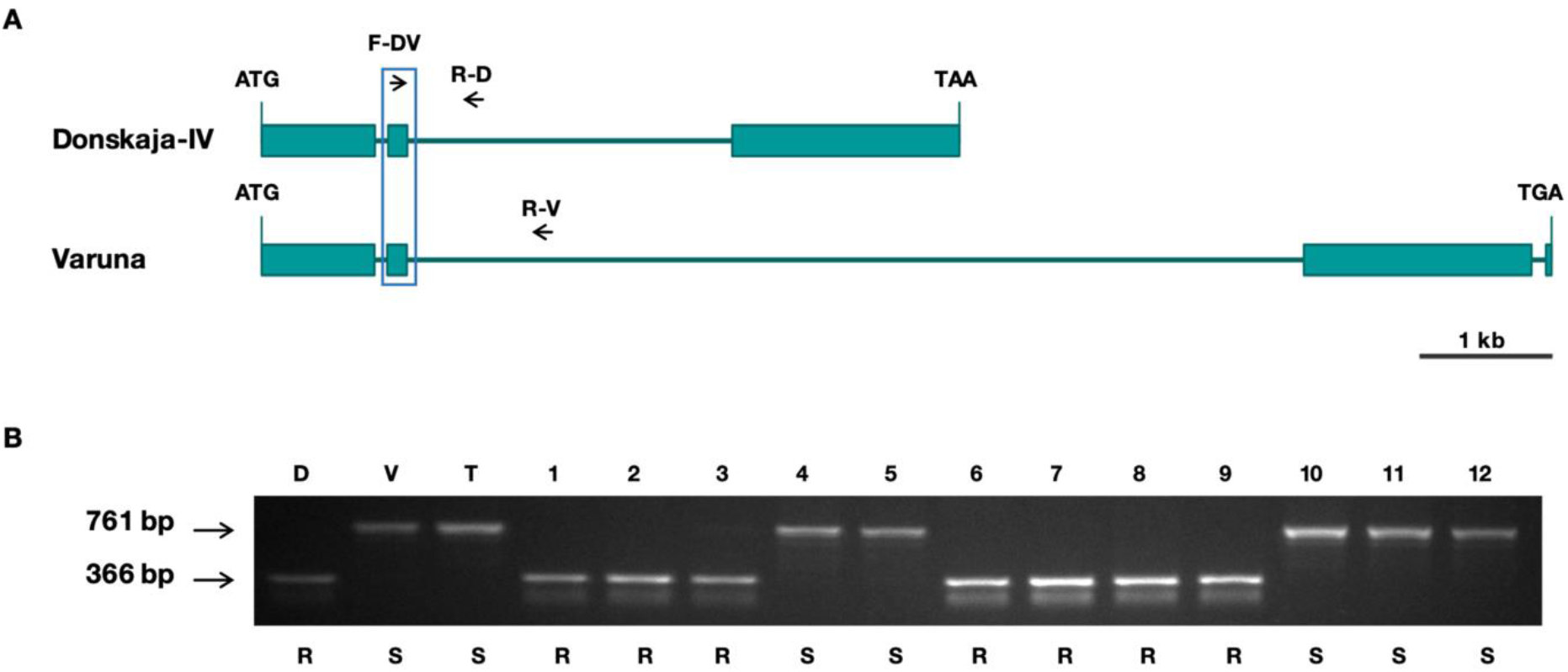
Development of *BjuWRR1* allele-specific marker and genotype analysis of TD doubled haploid (DH) mapping population. (A) Schematic representation of the sequences of *BjuWRR1* alleles of Donskaja-IV and Varuna. F-DV is a common forward primer whereas R-D and R-V are reverse primers specific to the Donskaja-IV and Varuna alleles, respectively. (B) Polymerase chain reaction (PCR) products amplified from genomic DNA prepared from Donskaja-IV (D), Varuna (V), TM-4 (T) and 12 individuals of TD-DH population. Lines resistant (R) and susceptible (S) to *A. candida* isolate AcB1 are mentioned.

To further validate the usefulness of the developed marker in marker-assisted selection (MAS), the individuals of the TD population previously used in mapping the white rust resistance locus were genotyped. The reported phenotypic data (Panjabi-Massand *et al.*, 2010) obtained by performing infection assays were compared with the genotypic data and the genotype was found to co-segregate with the phenotype (Fig. 8B). This marker can, therefore, be used for the introgression of Donskaja-IV_*BjuWRR1* in any of the Indian gene pool *B. juncea* lines.

## Discussion

Plants have evolved sophisticated defense mechanisms against pathogens besides the physical and chemical barriers. These mechanisms involve two layers of defense – PAMP-triggered immunity (PTI) is the first layer of defense that involves recognition of conserved pathogen-associated molecular patterns (PAMPs) of the invading pathogen via cell surface receptors (Pattern recognition receptors or PRR), and Effector-triggered immunity (ETI) is the second layer of defense involving recognition of the pathogen-secreted effectors (Avr) via intracellular receptors encoded by resistance (R) genes (Jones and Dangl, 2006; Dodds and Rathjen, 2010). Most plant R genes belong to the NB-LRR superfamily containing a Nucleotide-Binding (NB) domain followed by multiple Leucine-Rich Repeats (LRR) at the C-terminal end. NB-LRR proteins are subdivided into two major groups – TNL (TIR-NB-LRR) and CNL (CC-NB-LRR) on the basis of the presence of either a TIR (similar to animal Toll-like/interleukin-1 receptors) or a CC (coiled-coil) domain at the N-terminus (Michelmore *et al.*, 2013; Lee and Yeom, 2015).

In this study, we have identified *BjuWRR1*, a constitutively expressing R gene that encodes a CC-NB-LRR domain-containing protein, and provided a functional characterization of the gene and shown conclusively the involvement of this gene in conferring complete resistance to white rust in *B. juncea* line Donskaja-IV. R genes belonging to CNL class have been found to confer resistance to many pathogens in different crops and model species – RPS2, Rpm1 in Arabidopsis confer resistance against different pathovars of the bacterium, *Pseudomonas syringae* (Bent *et al.*, 1994; Mindrinos *et al.*, 1994; Grant *et al.*, 1995),Yr10 in wheat against fungus *Puccinia striiformis* (Liu *et al.*, 2014 a) and Pi64 in Rice against fungus *Magnaporthe oryzae* (Ma *et al.*, 2015). CNL type R genes conferring resistance to oomycete pathogens have also been described – RPP7 in Arabidopsis to *Hyaloperonospora arabidopsidis* (Eulgem *et al.*, 2007) and R3a (Huang *et al.*, 2005) and R3b (Li *et al.*, 2011) in Potato against *Phytophthora infestans*. However, to date, no CNL type gene conferring resistance to the oomycete pathogen *A. candida* has been reported.

The *BjuWRR1* gene from Donskaja-IV shows complete resistance to various isolates of *A. candida* as is evident from the analysis of transgenic lines developed using this gene in susceptible *B. juncea* line Varuna. *BjuWRR1*, therefore, has immense potential for developing *B. juncea* lines with complete and broad-spectrum white rust resistance. It would be interesting to assess the extent of resistance achieved by *BjuWRR1* against different races and isolates of *A. candida* collected from mustard growing regions other than India. The white rust resistance-conferring locus, AcB1-A5.1 harboring *BjuWRR1* has been introgressed into some of the extensively grown susceptible lines of the Indian gene pool of *B. juncea* namely, Varuna, Pusa bold, Pusa jai kisan and Rohini. These lines are ready to be released for cultivation. The locus can be further diversified into new lines and hybrids using the allele-specific molecular marker reported in this study.

Resistance conferred by the R gene against pathogens tend to break down. Therefore, more R genes will be required for long-term management of the white rust disease. One of the approaches to increase the durability of the resistance is to exploit the naturally occurring allelic variants and orthologues of a well-characterized R gene. Such an approach of allele mining has been deployed in wheat for managing powdery mildew wherein different alleles of the resistance gene *Pm3* have been cloned from different accessions of wheat and have been shown to confer resistance (Bhullar *et al.*, 2010). Functional orthologues of the gene *Pid3* that confer resistance to rice blast with different resistance spectra have been isolated and cloned (Shang *et al.*, 2009; Lv *et al.*, 2013; Xu *et al.*, 2014).

In the present work, we have studied the allelic variants and syntenic orthologues of *BjuWRR1* which could be tested to identify novel white rust resistant gene(s) with either same or different spectrum of resistance. Investigation of the functional and non-functional *BjuWRR1* alleles in Donskaja-IV, Cutlass, Varuna, Heera, Tumida, Skorospieka and Kranti suggests the presence of three types of alleles based on the sequence of the coding region. However, there was no correlation between the gene pool the line belonged to and the type of allele present. Varuna_*BjuWRR1* is non-functional for resistance to white rust since Varuna is highly susceptible to all the tested isolates. Alleles in Heera, Tumida, Skorospieka and Kranti having the identical coding sequence to the Varuna allele could also be non-functional in terms of conferring resistance to white rust. Non-functionality of *BjuWRR1* alleles in Heera and Tumida to white rust is also evident from the mapping of white rust resistance loci in Heera to the LG A04 (Panjabi-Massand *et al.*, 2010) and in Tumida to LG A06 (unpublished data). However, the transcription of the *BjuWRR1* alleles was not affected as we were able to detect transcripts of the gene in the tested lines – Varuna, Heera and Tumida. Cutlass, an east European gene pool line resistant to all the tested isolates, would be an interesting source to check whether Cutlass*_BjuWRR1* in spite of the truncation can confer resistance to white rust by developing transgenics in a susceptible *B. juncea* line like Varuna.

The search for *BjuWRR1* orthologues in different members of Brassicaceae family suggests that this CNL type R gene(s) is specific to the Brassica lineage. Analysis of the region syntenic to the locus AcB1-A5.1 in the Brassica A, B and C genomes, of diploid *B. rapa* (AA), *B. nigra* (BB) and *B. oleracea* (CC) and allotetraploid *B. juncea* (AABB) and *B. napus* (AACC) revealed the presence of at least one copy or remnants of CNL encoding R-genes in the syntenic regions suggesting that the ancestral CNL gene was present at this locus in the parent that contributed the LF genome to the mesohexaploid genomes of the three diploid species – *B. rapa*, *B. nigra* and *B. oleracea*. Differences in the haplotypes of the A subgenomes of the analyzed lines of *B. juncea* and *B. napus* suggests that different types of *B. rapa* have contributed their genomes to these allotetraploid species. We identified about eight genes from *B. rapa*, *B. napus* and *B. oleracea* that contain all the three domains of atypical CNL gene. Functional testing of these orthologues may lead to the identification of additional white rust resistance-conferring genes with either same or distinct resistance spectrum.

We conclude that the R gene *BjuWRR1* provides complete white rust resistance to all the tested *A. candida* isolates. This makes it an excellent resource for developing lines and hybrids resistant to the disease for the Indian subcontinent and sets the stage for a more methodical approach to managing white rust disease by studying the interaction between the allelic variants and orthologues of *BjuWRR1* from diverse Brassica germplasm with the Avr diversity present in the divergent isolates of *A. candida*.

## Supplementary data

**Fig. S1.** Schematic representation of the T-DNA of the binary vector used for the genetic transformation experiments.

**Fig. S2.** Schematic representation of the placement of the probes used for Southern hybridization analysis.

**Fig. S3.** Alignment of the nucleotide sequences of the coding region of *BjuWRR1* alleles and the amino acid sequences of the encoded proteins.

**Table S1.** List of germplasm used in the study.

**Table S2.** List of primers used in various experiments.

**Table S3.** Conserved domains predicted in the ORFs present in Donskaja-IV BAC containing the white rust resistance conferring locus AcB1-A5.1.

**Table S4.** Number of copies of the *bar* and *BjuWRR1* genes in different transgenic lines.

**Table S5.** Segregation for phosphinothricin resistance amongst progeny of the transgenic lines BjuWRR1-2 and BjuWRR1-3.

**Table S6.** Details of the genomes analyzed for *BjuWRR1* orthologues, the genes flanking the *BjuWRR1* orthologues and ID of the predicted CC-NB-LRR orthologues.

## Acknowledgments

We thank Mariam Ibrahim Taom for her help with cloning. This research was supported by the Department of Biotechnology (Government of India) with grants – BT/IN/Indo-UK/CGAT/12/DP/2014-15 and BT/01/NDDB/UDSC/2016. HA was supported by a research fellowship from UGC. DP was supported by a J. C. Bose Fellowship from the Department of Science & Technology (DST) and by the Council of Scientific and Industrial Research (CSIR) as a Distinguished Scientist.

## Figure legends

Colour in print: Fig. 1, Fig. 3, Fig. 6 and Fig. 7

Colour online-only: Fig. 2, Fig. 4, Fig. 5 and Fig. 8

