## Supplementary Data for "*BjuWRR1*, a CC-NB-LRR gene identified in *Brassica juncea*, confers resistance to white rust caused by *Albugo Candida*"

**Table S1.** List of germplasm used in the study.

| Germplasm | | Used as/ for | Origin |
|---|---|---|---|
| **Reference lines** | | | |
| *B. juncea* (AABB) | Donskaja-IV | Infection studies<br>Differential set<br>Gene isolation and cloning experiments<br>Expression studies | Russia |
| | Varuna | Infection studies<br>Differential set<br>Expression studies<br>Genetic transformation | India |
| | Heera | Differential set<br>Expression studies | Canada |
| | Tumida | Differential set<br>Expression studies | China |
| | Cutlass | Differential set | Poland |
| | Skorospieka | Differential set | Russia |
| *B. rapa* (AA) | YSPB | Differential set | India |
| | Chiifu-401 | Differential set | South Korea |
| | Candle | Differential set | Canada |
| | Pant toria | Differential set | India |
| *B. nigra* (BB) | Sangam | Differential set | India |
| | BN2782 | Differential set | |
| **Experimental lines** | | | |
| Near isogenic lines (Varuna) | Varuna-936-266 | Infection studies | |
| | Varuna-865-245 | Infection studies | |
| | Varuna-583-37 | Infection studies | |
| Near isogenic lines (Pusa bold) | PB-A5-223 | Infection studies | |
| Near isogenic lines (Pusa jai kisan) | PJK-A5-63 | Infection studies | |
| Near isogenic lines (Rohini) | Rohini-A5-467 | Infection studies | |
| Transgenic lines (Varuna) | BjuWRR1-2 | Infection studies | |
| | BjuWRR1-3 | Infection studies | |
| **TD-DH Mapping Population*** | | Genotyping study for validating allele specific markers | |

* Mapping population earlier used by Panjabi-Massand *et al.*, 2010

**Table S2.** List of primers used in various experiments.

| Primer | | Sequence (5′ to 3′) |
|---|---|---|
| At2g36360 | F | GCCACCTCCTAGATGTGGTCATA |
| | R | GTCCATCCAGGTGTTTCACG |
| At2g44970 | F | CCCACATTCCTTAAGCTCCAG |
| | R | CGGCTTCATTAAGAGCTGGT |
| At2g34700 | F | TACGCTCCTAGAAGCATCTCCTC |
| | R | CTCTCTTTGTGTTGTTGCATGC |
| At2g36480 | F | TTCATGCTGCAAAGGCCGTC |
| | R | TGCGGATGAAAGACTCCTTTCCAG |
| At2g36310 | F | CCTTCTCCTTGTAGAAAGAAATGTGAC |
| | R | CCAGAACACATGATGAATGTGACA |
| DIG-BjuWRR1 | F | CTCGACAGCAACTTGTGGAA |
| | R | TCCACCTTTTGAACACACCA |
| DIG-bar | F | ATGAGCCCAGAACGACG |
| | R | TACCGGCAGGCTGAAGTC |
| BjuWRR1inf | F | CCTCTAGAGTCGACCTGCAGGCATATACCAAGAGGCAAGAGTAGT |
| | R | AGCTCAAGCTAAGCTTTACAAGAAGAACAAGCCAAGACAG |
| UBQ9 | F | GAAGACATGTTCCATTGGCA |
| | R | ACACCTTAGTCCTAAAAGCCACCT |
| RT- BjuWRR1 | F | TTTGCATGTAAACCCAAAATC |
| | R | ATTTTCCTTCCCATGGCTTC |
| BjuWRR1seq | F | ATGGCTGACGGAGTTGTGTC |
| | R | TTATTCGTCGCTATCAGAGTCGTT |
| Bar | F | ACCATCGTCAACCACTACATC |
| | R | GTACCGGCAGGCTGAAGTC |
| BjuWRR1 | F | GGTTGTTAATCTGTCATACGCATC |
| | R | GCTCTCACCTTAAATGTTAAAATCGG |
| CDSE1-2 | F | CAGGCTAAGATCTTTCACTTGTTGG |
| | R | TGTTCGGTCTGCATGGAAAC |
| CDSE2-3 | F | GCTATGTGAGGGCATAGTATTCC |
| | R | GGCTAATGGTAGTCCTCCACAATA |
| CDSE3-4 | F | ACATAAGAAGAAGGAGTGGGAGAC |
| | R | TAGTGGTGGGTACTGATCCTGA |
| F-DV | F | GGCATAGTATTCCCTAGAAGAGAGATAAC |
| R-D | R | TGTTGATTCTTAGAATGGTAAATCACAG |
| R-V | R | TTGAAAATCACATGTATACATATGGCTT |

**Table S3.** Conserved domains predicted in the ORFs present in Donskaja-IV BAC containing the white rust resistance conferring locus AcB1-A5.1. The domains were predicted using an online tool – NCBI Batch CD-Search.

| Query | Hit type | Accession | Superfamily | Short name | Definition |
|---|---|---|---|---|---|
| ORF1 | specific | pfam00046 | cl00084 | Homeobox | Homeobox domain |
| | superfamily | cl23880 | - | HALZ superfamily | Homeobox associated leucine zipper |
| ORF2 | specific | pfam03031 | cl21460 | NIF | NLI interacting factor-like phosphatase |
| | superfamily | cl21460 | - | HAD_like superfamily | Haloacid Dehalogenase-like Hydrolases |
| ORF3 | superfamily | cl12382 | | Rogdi_lz superfamily | Rogdi leucine zipper containing protein |
| ORF4 | superfamily | cl21453 | | PKc_like superfamily | Protein Kinases |
| ORF5 | specific | PLN00191 | cl27211 | PLN00191 | Enolase |
| | superfamily | cl21457 | - | ICL_KPHMT superfamily | Members of the ICL/PEPM_KPHMT enzyme superfamily |
| ORF6 | specific | pfam09733 | cl10718 | VEFS-Box | VEFS-Box of polycomb protein |
| ORF7 | specific | cd04587 | cl15354 | CBS_pair_CAP-ED_DUF294_PBI_assoc | CBS domain associated with CAP_ED and DUF294 domain or PB1 domain |
| | specific | pfam00564 | cl02720 | PB1 | PB1 domain |
| | specific | COG0517 | cl15354 | CBS | CBS domain |
| | superfamily | cl27608 | - | IMPDH superfamily | IMP dehydrogenase / GMP reductase domain |
| | superfamily | cl25376 | - | gutQ superfamily | D-arabinose 5-phosphate isomerase |
| | superfamily | cl27346 | - | DHHA2 superfamily | DHHA2 domain |
| | superfamily | cl27264 | - | CorC_HlyC superfamily | Transporter associated domain |
| ORF8 | specific | pfam03004 | cl03830 | Transposase_24 | Plant transposase (Ptta/En/Spm family) |
| ORF9 | specific | pfam15628 | cl21422 | RRM_DME | RRM in Demeter |
| | specific | cd03562 | cl02544 | CID | CID (CTD-Interacting Domain) domain family |
| | superfamily | cl02544 | - | VHS_ENTH_ANTH superfamily | VHS, ENTH and ANTH domain superfamily |
| | specific | smart00582 | cl02544 | RPR | Domain present in proteins involved in regulation of nuclear pre-mRNA |
| | superfamily | cl27658 | - | HHH superfamily | Helix-hairpin-helix motif |
| | specific | smart00478 | cl23768 | ENDO3c | Endonuclease III |
| | specific | pfam15629 | cl21423 | Perm-CXXC | Permuted single zf-CXXC unit |
| | specific | pfam04818 | cl02544 | CTD_bind | RNA polymerase II-binding domain |
| | superfamily | cl28533 | - | ogg superfamily | 8-oxoguanine DNA-glycosylase (ogg) |
| | superfamily | cl26830 | - | AlkA superfamily | 3-methyladenine DNA glycosylase/8-oxoguanine DNA glycosylase |
| | superfamily | cl26512 | - | EndIII_4Fe-2S superfamily | Iron-sulfur binding domain of endonuclease III |
| | superfamily | cl00368 | - | Ribosomal_S16 superfamily | Ribosomal protein S16 |
| ORF10 | specific | pfam05910 | cl05466 | DUF868 | Plant protein of unknown function |
| ORF11 | superfamily | cl25404 | - | GH38-57_N_LamB_YdjC_SF superfamily | Domain of glycoside hydrolase, lactam utilization protein, YdjC-family protein |
| ORF12 | specific | pfam03140 | cl03911 | DUF247 | Plant protein of unknown function |
| ORF13 | superfamily | cl16789 | - | DUF4378 superfamily | Domain of unknown function |
| ORF14 | superfamily | cl02808 | - | RT_like superfamily | Reverse transcriptase_like family |
| | specific | pfam13966 | cl16506 | zf-RVT | Zinc-binding in reverse transcriptase |
| ORF15 | superfamily | cl27543 | - | PRK13004 superfamily | Peptidase; Reviewed |

| | | | | | |
|---|---|---|---|---|---|
| ORF16 | superfamily | cl27380 | - | Ald_Xan_dh_C2 superfamily | Molybdopterin-binding domain of aldehyde dehydrogenase |
| ORF17 | specific | cd01650 | cl02808 | RT_nLTR_like | Non-LTR retrotransposon and non-LTR retrovirus reverse transcriptase |
| | specific | pfam03372 | cl00490 | Exo_endo_phos | Endonuclease/Exonuclease/phosphatase family |
| | superfamily | cl28246 | - | DnaJ superfamily | DnaJ-class molecular chaperone with Zn finger domain |
| | superfamily | cl26511 | - | Neuromodulin_N superfamily | Gap junction protein N-terminal region |
| | superfamily | cl22451 | - | ASF1_hist_chap superfamily | ASF1 like histone chaperone |
| | superfamily | cl24027 | - | SDA1 superfamily | Consists of SDA1 protein homologs |
| | superfamily | cl27511 | - | Na_Ca_ex superfamily | Sodium/calcium exchanger protein |
| | superfamily | cl03959 | - | Prothymosin superfamily | Prothymosin/parathymosin family |
| ORF18 | superfamily | cl28391 | - | PurB superfamily | Adenylosuccinate lyase |
| ORF19 | specific | pfam00665 | cl21549 | rve | Integrase core domain |
| | specific | pfam14244 | cl28789 | Retrotran_gag_3 | Gag-polypeptide of LTR copia-type |
| ORF20 | specific | pfam08879 | cl07468 | WRC | WRC domain |
| | specific | pfam08880 | cl07469 | QLQ | QLQ domain |
| ORF21 | specific | PLN02447 | cl25948 | PLN02447 | 1,4-alpha-glucan-branching enzyme |
| | superfamily | cl25948 | - | PulA superfamily | Pullulanase/glycogen debranching enzyme |
| | specific | cd11321 | cl07893 | AmyAc_bac_euk_BE | Alpha amylase catalytic domain |
| ORF22 | specific | pfam00931 | cl26397 | NB-ARC | NB-ARC domain |
| | specific | cd14798 | cl26396 | RX-CC_like | Coiled-coil domain of the potato virux X and similar resistance protein |
| | superfamily | cl27891 | - | LRR_3 superfamily | Leucine Rich Repeat |
| | superfamily | cl26793 | - | PLN00113 superfamily | Leucine-rich repeat receptor-like protein kinase; Provisional |
| | superfamily | cl21455 | - | P-loop_NTPase superfamily | P-loop containing Nucleoside Triphosphate Hydrolases |
| ORF23 | superfamily | cl26061 | - | PLN02193 superfamily | Nitrile-specifier protein |
| | specific | pfam13415 | cl02701 | Kelch_3 | Galactose oxidase, central domain |
| | superfamily | cl27059 | - | Guanylate_kin superfamily | Guanylate kinase |
| | superfamily | cl28748 | - | RAG2 superfamily | Recombination activating protein 2 |
| | specific | smart00612 | cl02701 | Kelch | Kelch domain |
| ORF24 | specific | smart00220 | cl26011 | S_TKc | Serine/Threonine protein kinases |
| | superfamily | cl27677 | - | PRK07788 superfamily | Acyl-CoA synthetase; Validated |
| ORF25 | specific | pfam04535 | cl04571 | DUF588 | Domain of unknown function |
| | superfamily | cl04571 | - | MARVEL superfamily | Membrane-associating domain |
| ORF26 | specific | smart00154 | cl01438 | ZnF_AN1 | AN1-like Zinc finger |
| | specific | pfam01754 | cl27506 | zf-A20 | A20-like zinc finger |
| ORF27 | specific | cd04476 | cl09930 | RPA1_DBD_C | RPA1_DBD_C |
| | specific | cd04480 | cl09930 | RPA1_DBD_A_like | RPA1_DBD_A_like |
| | superfamily | cl28407 | - | RFA1 superfamily | ssDNA-binding replication factor A, large subunit |
| ORF28 | specific | PLN02717 | cl00226 | PLN02717 | Uridine nucleosidase |

**Table S4.** Number of copies of the *bar* and *BjuWRR1* genes in different transgenic lines. Number was determined by Southern hybridization using *bar* and *BjuWRR1* gene specific probes.

| **Transgenic lines** | **Number of copies** | |
|---|---|---|
| | ***bar*** | ***BjuWRR1*** |
| BjuWRR1-1 | 3 | 3 |
| BjuWRR1-2 | 1 | 1 |
| BjuWRR1-3 | 1 | 1 |
| BjuWRR1-4 | 3 | 3 |
| BjuWRR1-6 | 3 | 2 |
| BjuWRR1-7 | 3 | 3 |
| BjuWRR1-10 | 2 | 2 |
| BjuWRR1-11 | 2 | 2 |
| BjuWRR1-13 | 3 | 2 |
| BjuWRR1-14 | 3 | 2 |
| BjuWRR1-15 | 3 | 1 |
| BjuWRR1-16 | 3 | 1 |
| BjuWRR1-18 | 4 | 3 |
| BjuWRR1-19 | 8 | 6 |
| BjuWRR1-21 | 1 | 1 |
| BjuWRR1-25 | 2 | 1 |
| BjuWRR1-27 | 2 | 1 |
| BjuWRR1-28 | 2 | 2 |
| BjuWRR1-29 | 2 | 2 |
| BjuWRR1-32 | 1 | 1 |
| BjuWRR1-33 | 1 | 1 |
| BjuWRR1-34 | 1 | 1 |
| BjuWRR1-38 | 1 | 4 |
| BjuWRR1-39 | 1 | 2 |
| BjuWRR1-40 | 1 | 3 |
| BjuWRR1-42 | 3 | 3 |
| BjuWRR1-43 | 2 | 2 |
| BjuWRR1-44 | 2 | 5 |

**Table S5.** Segregation for phosphinothricin resistance amongst progeny of the transgenic lines BjuWRR1-2 and BjuWRR1-3.

| **Transgenic line** | **Total $T_1$** | **R** | **S** | $\chi^2$ | **Number of copies** | |
|---|---|---|---|---|---|---|
| | | | | | ***bar*** | ***BjuWRR1*** |
| BjuWRR1-2 | 31 | 22 | 9 | 0.267 | 1 | 1 |
| BjuWRR1-3 | 43 | 30 | 13 | 0.626 | 1 | 1 |

$T_1$ seedlings were assessed for resistance (R) or susceptibility (S) to PPT (200 mg $L^{-1}$); chi-square was performed at 95% confidence limit ($p < 0.05$) to determine the goodness-of-fit. The data fits into a 3:1 ratio for resistance and susceptibility to PPT.

**Table S6.** Details of the genomes analyzed for *BjuWRR1* orthologues, the genes flanking the *BjuWRR1* orthologues and ID of the predicted CC-NB-LRR orthologues.

| Genome | Variety | Chromosome | Flanking Genes | CNL Genes | Sequence source |
|---|---|---|---|---|---|
| *A. thaliana* | Columbia (Col-0) | Chr. 2 | At2g36390 – At2g36350 | - | BRAD |
| *B. rapa* (AA) | Chiifu-401 | A05 | Bra005269 – Bra005280 | Bra005270*, Bra005271, Bra005273, Bra005277*, Bra005278* | BRAD |
| *B. oleracea* (CC) | Capitata line 02-12 | C04 | Bol011731 – Bol011727 | Bol011729* | BRAD |
| *B. napus* (AACC) | Darmor-bzh | A05, C04 | AA: GSBRNA2T00117069001 – GSBRNA2T00117081001<br>CC: GSBRNA2T00141135001 – GSBRNA2T00141142001 | AA: GSBRNA2T00117070001, GSBRNA2T00117071001, GSBRNA2T00117074001, GSBRNA2T00117079001<br>CC: GSBRNA2T00141140001 | BRAD |
| *B. nigra* (BB) | Sangam | B04 | B04_Sangam_g4068 – B04_Sangam_g4066 | Remnants | In house |
| *B. juncea* (AABB) | Donskaja-IV | A05, B04 | AA: Bju_A05_g947 – Bju_A05_g949<br>BB: Bju_B04_g11063 – Bju_B04_g11061 | AA: Bju_A05_g948A<br>BB: Remnants | In house |

*Genes lacking CC, NB and/or LRR domains

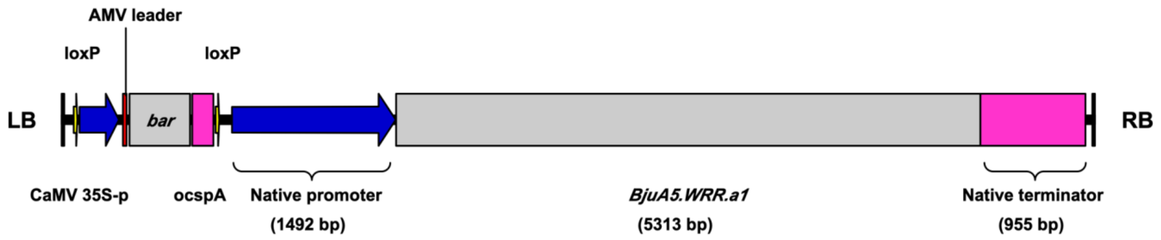


**Fig. S1.** Schematic representation of the T-DNA of the binary vector used for the genetic transformation experiments. Genomic regions of the *bar* and *BjuA5.WRR.a1* genes from start to stop codons, are shown as grey boxes. The *bar* gene is driven by CaMV35S promoter and an AMV leader sequence and has ocspA as the terminator sequence and was cloned between the two loxP sites. Blue and pink boxes represent the promoter and terminator regions, respectively. LB and RB stands for left and right border of the T-DNA.

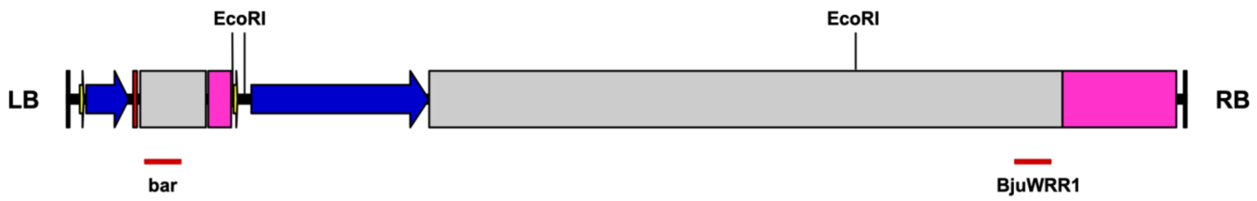


**Fig. S2.** Schematic representation of the placement of the probes used for Southern hybridization analysis. The position of the three EcoRI sites and the *bar* and *BjuWRR1* gene specific probes are highlighted.

10 20 30 40 50

| | | | | | | | | | | | | | | | | | | |
|---|---|---|---|---|---|---|---|---|---|---|---|---|---|---|---|---|---|---|
| Donskaja-IV | M | A | D | G | V | V | S | F | G | V | E | K | L | W | D | L | L | S |
| | ATG | GCT | GAC | GGA | GTT | GTG | TCG | TTT | GGA | GTG | GAG | AAA | CTC | TGG | GAT | CTC | CTG | AGT |
| Cutlass | M | A | D | G | V | V | S | F | G | V | E | K | L | W | D | L | L | S |
| | ATG | GCT | GAC | GGA | GTT | GTG | TCG | TTT | GGA | GTG | GAG | AAA | CTC | TGG | GAT | CTC | CTG | AGT |
| Varuna | M | A | E | G | V | V | S | F | G | V | E | K | L | W | D | L | L | S |
| | ATG | GCT | GAG | GGC | GTT | GTG | TCG | TTT | GGA | GTG | GAG | AAA | CTC | TGG | GAT | CTC | CTG | AGT |
| Heera | M | A | E | G | V | V | S | F | G | V | E | K | L | W | D | L | L | S |
| | ATG | GCT | GAG | GGC | GTT | GTG | TCG | TTT | GGA | GTG | GAG | AAA | CTC | TGG | GAT | CTC | CTG | AGT |
| Skorospieka | M | A | E | G | V | V | S | F | G | V | E | K | L | W | D | L | L | S |
| | ATG | GCT | GAG | GGC | GTT | GTG | TCG | TTT | GGA | GTG | GAG | AAA | CTC | TGG | GAT | CTC | CTG | AGT |
| Kranti | M | A | E | G | V | V | S | F | G | V | E | K | L | W | D | L | L | S |
| | ATG | GCT | GAG | GGC | GTT | GTG | TCG | TTT | GGA | GTG | GAG | AAA | CTC | TGG | GAT | CTC | CTG | AGT |
| Tumida | M | A | E | G | V | V | S | F | G | V | E | K | L | W | D | L | L | S |
| | ATG | GCT | GAG | GGC | GTT | GTG | TCG | TTT | GGA | GTG | GAG | AAA | CTC | TGG | GAT | CTC | CTG | AGT |

60 70 80 90 100

| | | | | | | | | | | | | | | | | | | |
|---|---|---|---|---|---|---|---|---|---|---|---|---|---|---|---|---|---|---|
| Donskaja-IV | R | E | S | E | R | L | Q | G | V | H | E | K | V | D | D | L | K | C |
| | CGA | GAA | TCA | GAG | AGA | CTG | CAG | GGA | GTT | CAT | GAG | AAA | GTT | GAT | GAT | CTA | AAA | TGT |
| Cutlass | R | E | S | E | R | L | Q | G | V | H | E | K | V | D | D | L | K | C |
| | CGA | GAA | TCA | GAG | AGA | CTG | CAG | GGA | GTT | CAT | GAG | AAA | GTT | GAT | GAT | CTA | AAA | TGT |
| Varuna | R | E | S | E | R | L | Q | G | V | H | E | K | V | D | D | L | K | C |
| | CGA | GAA | TCT | GAG | AGA | CTG | CAG | GGA | GTT | CAT | GAG | AAA | GTT | GAT | GAT | CTA | AAA | TGT |
| Heera | R | E | S | E | R | L | Q | G | V | H | E | K | V | D | D | L | K | C |
| | CGA | GAA | TCT | GAG | AGA | CTG | CAG | GGA | GTT | CAT | GAG | AAA | GTT | GAT | GAT | CTA | AAA | TGT |
| Skorospieka | R | E | S | E | R | L | Q | G | V | H | E | K | V | D | D | L | K | C |
| | CGA | GAA | TCT | GAG | AGA | CTG | CAG | GGA | GTT | CAT | GAG | AAA | GTT | GAT | GAT | CTA | AAA | TGT |
| Kranti | R | E | S | E | R | L | Q | G | V | H | E | K | V | D | D | L | K | C |
| | CGA | GAA | TCT | GAG | AGA | CTG | CAG | GGA | GTT | CAT | GAG | AAA | GTT | GAT | GAT | CTA | AAA | TGT |
| Tumida | R | E | S | E | R | L | Q | G | V | H | E | K | V | D | D | L | K | C |
| | CGA | GAA | TCT | GAG | AGA | CTG | CAG | GGA | GTT | CAT | GAG | AAA | GTT | GAT | GAT | CTA | AAA | TGT |

110 120 130 140 150 160

| | | | | | | | | | | | | | | | | | | |
|---|---|---|---|---|---|---|---|---|---|---|---|---|---|---|---|---|---|---|
| Donskaja-IV | Q | M | R | M | L | Q | S | L | L | K | D | A | D | A | R | K | Y | E |
| | CAG | ATG | AGA | ATG | TTA | CAG | TCG | TTG | TTG | AAA | GAT | GCA | GAT | GCC | AGG | AAA | TAT | GAG |
| Cutlass | Q | M | R | M | L | Q | S | L | L | K | D | A | D | A | R | K | Y | E |
| | CAG | ATG | AGA | ATG | TTA | CAG | TCG | TTG | TTG | AAA | GAT | GCA | GAT | GCC | AGG | AAA | TAT | GAG |
| Varuna | Q | M | R | M | L | Q | S | L | L | K | D | A | D | A | K | K | Y | E |
| | CAG | ATG | AGA | ATG | TTA | CAG | TCG | TTG | TTG | AAA | GAT | GCA | GAT | GCC | AAG | AAA | TAT | GAG |
| Heera | Q | M | R | M | L | Q | S | L | L | K | D | A | D | A | K | K | Y | E |
| | CAG | ATG | AGA | ATG | TTA | CAG | TCG | TTG | TTG | AAA | GAT | GCA | GAT | GCC | AAG | AAA | TAT | GAG |
| Skorospieka | Q | M | R | M | L | Q | S | L | L | K | D | A | D | A | K | K | Y | E |
| | CAG | ATG | AGA | ATG | TTA | CAG | TCG | TTG | TTG | AAA | GAT | GCA | GAT | GCC | AAG | AAA | TAT | GAG |
| Kranti | Q | M | R | M | L | Q | S | L | L | K | D | A | D | A | K | K | Y | E |
| | CAG | ATG | AGA | ATG | TTA | CAG | TCG | TTG | TTG | AAA | GAT | GCA | GAT | GCC | AAG | AAA | TAT | GAG |
| Tumida | Q | M | R | M | L | Q | S | L | L | K | D | A | D | A | K | K | Y | E |
| | CAG | ATG | AGA | ATG | TTA | CAG | TCG | TTG | TTG | AAA | GAT | GCA | GAT | GCC | AAG | AAA | TAT | GAG |

170 180 190 200 210

| | | | | | | | | | | | | | | | | | | |
|---|---|---|---|---|---|---|---|---|---|---|---|---|---|---|---|---|---|---|
| Donskaja-IV | N | E | A | V | R | N | F | L | E | D | V | K | D | I | V | F | D | A |
| | AAT | GAA | GCA | GTG | AGA | AAC | TTC | TTG | GAA | GAT | GTC | AAA | GAC | ATT | GTA | TTT | GAT | GCT |
| Cutlass | N | - | A | V | R | N | F | L | E | D | V | K | D | I | V | F | D | A |
| | AAT | - - - | GCA | GTG | AGA | AAC | TTC | TTG | GAA | GAT | GTC | AAA | GAC | ATT | GTA | TTT | GAT | GCT |
| Varuna | S | - | V | V | R | N | F | L | E | D | V | K | D | T | V | F | D | A |
| | AGT | - - - | GTG | GTG | AGA | AAC | TTC | TTG | GAA | GAT | GTC | AAA | GAC | ACT | GTG | TTT | GAT | GCT |
| Heera | S | - | V | V | R | N | F | L | E | D | V | K | D | T | V | F | D | A |
| | AGT | - - - | GTG | GTG | AGA | AAC | TTC | TTG | GAA | GAT | GTC | AAA | GAC | ACT | GTG | TTT | GAT | GCT |
| Skorospieka | S | - | V | V | R | N | F | L | E | D | V | K | D | T | V | F | D | A |
| | AGT | - - - | GTG | GTG | AGA | AAC | TTC | TTG | GAA | GAT | GTC | AAA | GAC | ACT | GTG | TTT | GAT | GCT |
| Kranti | S | - | V | V | R | N | F | L | E | D | V | K | D | T | V | F | D | A |
| | AGT | - - - | GTG | GTG | AGA | AAC | TTC | TTG | GAA | GAT | GTC | AAA | GAC | ACT | GTG | TTT | GAT | GCT |
| Tumida | S | - | V | V | R | N | F | L | E | D | V | K | D | T | V | F | D | A |
| | AGT | - - - | GTG | GTG | AGA | AAC | TTC | TTG | GAA | GAT | GTC | AAA | GAC | ACT | GTG | TTT | GAT | GCT |

220 230 240 250 260 270

| | | | | | | | | | | | | | | | | | | |
|---|---|---|---|---|---|---|---|---|---|---|---|---|---|---|---|---|---|---|
| Donskaja-IV | E | D | I | I | E | S | F | L | L | K | E | L | S | G | N | Q | K | G |
| | GAA | GAT | ATA | ATT | GAA | TCC | TTT | CTT | TTG | AAA | GAA | CTT | AGT | GGA | AAC | CAA | AAA | GGG |
| Cutlass | E | D | I | I | E | S | F | L | L | K | E | L | S | G | N | Q | K | G |
| | GAA | GAT | ATA | ATT | GAA | TCC | TTT | CTT | TTG | AAA | GAA | CTT | AGT | GGA | AAC | CAA | AAA | GGG |
| Varuna | E | D | I | I | E | S | F | L | L | K | E | L | S | G | N | Q | K | G |
| | GAA | GAC | ATA | ATC | GAA | TCC | TTT | CTT | TTG | AAA | GAA | TTG | AGT | GGA | AAC | CAA | AAA | GGG |
| Heera | E | D | I | I | E | S | F | L | L | K | E | L | S | G | N | Q | K | G |
| | GAA | GAC | ATA | ATC | GAA | TCC | TTT | CTT | TTG | AAA | GAA | TTG | AGT | GGA | AAC | CAA | AAA | GGG |
| Skorospieka | E | D | I | I | E | S | F | L | L | K | E | L | S | G | N | Q | K | G |
| | GAA | GAC | ATA | ATC | GAA | TCC | TTT | CTT | TTG | AAA | GAA | TTG | AGT | GGA | AAC | CAA | AAA | GGG |
| Kranti | E | D | I | I | E | S | F | L | L | K | E | L | S | G | N | Q | K | G |
| | GAA | GAC | ATA | ATC | GAA | TCC | TTT | CTT | TTG | AAA | GAA | TTG | AGT | GGA | AAC | CAA | AAA | GGG |
| Tumida | E | D | I | I | E | S | F | L | L | K | E | L | S | G | N | Q | K | G |
| | GAA | GAC | ATA | ATC | GAA | TCC | TTT | CTT | TTG | AAA | GAA | TTG | AGT | GGA | AAC | CAA | AAA | GGG |

280 290 300 310 320

| | | | | | | | | | | | | | | | | | | |
|---|---|---|---|---|---|---|---|---|---|---|---|---|---|---|---|---|---|---|
| Donskaja-IV | I | K | G | R | V | K | R | L | S | C | F | L | V | D | R | R | G | L |
| | ATC | AAG | GGG | CGT | GTG | AAA | AGA | CTT | TCT | TGT | TTC | TTA | GTG | GAT | CGC | CGA | GGT | CTT |
| Cutlass | I | K | G | R | V | K | R | L | S | C | F | L | V | D | R | R | G | L |
| | ATC | AAG | GGG | CGT | GTG | AAA | AGA | CTT | TCT | TGT | TTC | TTA | GTG | GAT | CGC | CGA | GGT | CTT |
| Varuna | I | K | G | R | V | K | R | L | S | C | F | L | V | D | R | R | G | L |
| | ATC | AAG | GGG | CGT | GTG | AAA | AGA | CTT | TCT | TGT | TTC | TTA | GTG | GAT | CGC | CGA | GGT | CTT |
| Heera | I | K | G | R | V | K | R | L | S | C | F | L | V | D | R | R | G | L |
| | ATC | AAG | GGG | CGT | GTG | AAA | AGA | CTT | TCT | TGT | TTC | TTA | GTG | GAT | CGC | CGA | GGT | CTT |
| Skorospieka | I | K | G | R | V | K | R | L | S | C | F | L | V | D | R | R | G | L |
| | ATC | AAG | GGG | CGT | GTG | AAA | AGA | CTT | TCT | TGT | TTC | TTA | GTG | GAT | CGC | CGA | GGT | CTT |
| Kranti | I | K | G | R | V | K | R | L | S | C | F | L | V | D | R | R | G | L |
| | ATC | AAG | GGG | CGT | GTG | AAA | AGA | CTT | TCT | TGT | TTC | TTA | GTG | GAT | CGC | CGA | GGT | CTT |
| Tumida | I | K | G | R | V | K | R | L | S | C | F | L | V | D | R | R | G | L |
| | ATC | AAG | GGG | CGT | GTG | AAA | AGA | CTT | TCT | TGT | TTC | TTA | GTG | GAT | CGC | CGA | GGT | CTT |

```
              330 340 350 360 370
Donskaja-IV    S   M   D   I   E   G   I   T   K   R   M   S   K   V   V   A   Q   M
              TCT ATG GAC ATT GAA GGC ATA ACT AAG AGA ATG TCT AAG GTG GTT GCG CAA ATG
Cutlass        S   M   D   I   E   G   I   T   K   R   M   S   K   V   V   A   Q   M
              TCT ATG GAC ATT GAA GGC ATA ACT AAG AGA ATG TCT AAG GTG GTT GCG CAA ATG
Varuna         S   M   D   I   E   G   I   T   K   R   M   A   K   V   V   G   Q   M
              TCT ATG GAC ATT GAA GGC ATA ACT AAG AGA ATG GCC AAG GTT GTT GGG CAA ATG
Heera          S   M   D   I   E   G   I   T   K   R   M   A   K   V   V   G   Q   M
              TCT ATG GAC ATT GAA GGC ATA ACT AAG AGA ATG GCC AAG GTT GTT GGG CAA ATG
Skorospieka    S   M   D   I   E   G   I   T   K   R   M   A   K   V   V   G   Q   M
              TCT ATG GAC ATT GAA GGC ATA ACT AAG AGA ATG GCC AAG GTT GTT GGG CAA ATG
Kranti         S   M   D   I   E   G   I   T   K   R   M   A   K   V   V   G   Q   M
              TCT ATG GAC ATT GAA GGC ATA ACT AAG AGA ATG GCC AAG GTT GTT GGG CAA ATG
Tumida         S   M   D   I   E   G   I   T   K   R   M   A   K   V   V   G   Q   M
              TCT ATG GAC ATT GAA GGC ATA ACT AAG AGA ATG GCC AAG GTT GTT GGG CAA ATG

              380 390 400 410 420 430
Donskaja-IV    Q   S   F   G   I   Q   H   S   V   N   K   P   P   P   L   Q   K   R
              CAG AGC TTT GGT ATA CAG CAT AGC GTT AAC AAA CCA CCG CCT CTA CAA AAG AGA
Cutlass        Q   S   F   G   I   Q   H   S   V   N   K   P   P   P   L   Q   K   R
              CAG AGC TTT GGT ATA CAG CAT AGC GTT AAC AAA CCA CCG CCT CTA CAA AAG AGA
Varuna         Q   S   F   G   I   Q   Q   I   I   Y   R   S   P   S   L   Q   E   R
              CAG AGC TTT GGA ATA CAA CAG ATC ATA TAC AGA TCA CCG TCT CTA CAA GAG AGA
Heera          Q   S   F   G   I   Q   Q   I   I   Y   R   S   P   S   L   Q   E   R
              CAG AGC TTT GGA ATA CAA CAG ATC ATA TAC AGA TCA CCG TCT CTA CAA GAG AGA
Skorospieka    Q   S   F   G   I   Q   Q   I   I   Y   R   S   P   S   L   Q   E   R
              CAG AGC TTT GGA ATA CAA CAG ATC ATA TAC AGA TCA CCG TCT CTA CAA GAG AGA
Kranti         Q   S   F   G   I   Q   Q   I   I   Y   R   S   P   S   L   Q   E   R
              CAG AGC TTT GGA ATA CAA CAG ATC ATA TAC AGA TCA CCG TCT CTA CAA GAG AGA
Tumida         Q   S   F   G   I   Q   Q   I   I   Y   R   S   P   S   L   Q   E   R
              CAG AGC TTT GGA ATA CAA CAG ATC ATA TAC AGA TCA CCG TCT CTA CAA GAG AGA

              440 450 460 470 480
Donskaja-IV    Q   R   E   I   Q   Q   T   Y   P   K   R   S   Q   K   D   L   V   G
              CAG AGG GAG ATC CAA CAA ACA TAT CCA AAG AGA TCT CAA AAG GAT CTT GTG GGG
Cutlass        Q   R   E   I   Q   Q   T   Y   P   K   R   S   Q   K   D   L   V   G
              CAG AGG GAG ATC CAA CAA ACA TAT CCA AAG AGA TCT CAA AAG GAT CTT GTG GGG
Varuna         Q   R   E   I   Q   Q   T   Y   P   M   G   S   Q   K   D   L   V   G
              CAG AGG GAG ATC CAA CAA ACA TAT CCT ATG GGA TCT CAA AAG GAT CTT GTG GGG
Heera          Q   R   E   I   Q   Q   T   Y   P   M   G   S   Q   K   D   L   V   G
              CAG AGG GAG ATC CAA CAA ACA TAT CCT ATG GGA TCT CAA AAG GAT CTT GTG GGG
Skorospieka    Q   R   E   I   Q   Q   T   Y   P   M   G   S   Q   K   D   L   V   G
              CAG AGG GAG ATC CAA CAA ACA TAT CCT ATG GGA TCT CAA AAG GAT CTT GTG GGG
Kranti         Q   R   E   I   Q   Q   T   Y   P   M   G   S   Q   K   D   L   V   G
              CAG AGG GAG ATC CAA CAA ACA TAT CCT ATG GGA TCT CAA AAG GAT CTT GTG GGG
Tumida         Q   R   E   I   Q   Q   T   Y   P   M   G   S   Q   K   D   L   V   G
              CAG AGG GAG ATC CAA CAA ACA TAT CCT ATG GGA TCT CAA AAG GAT CTT GTG GGG

              490 500 510 520 530 540
Donskaja-IV    V   E   Q   S   V   N   E   L   V   G   H   L   V   E   N   D   N   I
              GTG GAA CAG AGT GTT AAT GAA TTG GTT GGT CAT TTG GTG GAG AAT GAT AAC ATC
Cutlass        V   E   Q   S   V   N   E   L   V   G   H   L   V   E   N   D   N   I
              GTG GAA CAG AGT GTT AAT GAA TTG GTT GGT CAT TTG GTG GAG AAT GAT AAC ATC
Varuna         V   E   Q   S   V   K   E   L   V   G   H   L   V   E   N   D   N   I
              GTG GAA CAG AGT GTT AAA GAA TTG GTT GGT CAT TTG GTG GAG AAT GAT AAC ATC
Heera          V   E   Q   S   V   K   E   L   V   G   H   L   V   E   N   D   N   I
              GTG GAA CAG AGT GTT AAA GAA TTG GTT GGT CAT TTG GTG GAG AAT GAT AAC ATC
Skorospieka    V   E   Q   S   V   K   E   L   V   G   H   L   V   E   N   D   N   I
              GTG GAA CAG AGT GTT AAA GAA TTG GTT GGT CAT TTG GTG GAG AAT GAT AAC ATC
Kranti         V   E   Q   S   V   K   E   L   V   G   H   L   V   E   N   D   N   I
              GTG GAA CAG AGT GTT AAA GAA TTG GTT GGT CAT TTG GTG GAG AAT GAT AAC ATC
Tumida         V   E   Q   S   V   K   E   L   V   G   H   L   V   E   N   D   N   I
              GTG GAA CAG AGT GTT AAA GAA TTG GTT GGT CAT TTG GTG GAG AAT GAT AAC ATC

              550 560 570 580 590
Donskaja-IV    Q   V   V   S   I   S   G   M   G   G   I   G   K   T   T   L   A   R
              CAA GTA GTT TCC ATA TCT GGG ATG GGT GGT ATT GGG AAA ACC ACT CTT GCC AGA
Cutlass        Q   V   V   S   I   S   G   M   G   G   I   G   K   T   T   L   A   R
              CAA GTA GTT TCC ATA TCT GGG ATG GGT GGT ATT GGG AAA ACC ACT CTT GCC AGA
Varuna         Q   V   V   S   I   S   G   M   G   G   I   G   K   T   T   L   A   R
              CAA GTA GTT TCC ATA TCT GGG ATG GGT GGT ATT GGA AAA ACC ACT CTT GCC AGA
Heera          Q   V   V   S   I   S   G   M   G   G   I   G   K   T   T   L   A   R
              CAA GTA GTT TCC ATA TCT GGG ATG GGT GGT ATT GGA AAA ACC ACT CTT GCC AGA
Skorospieka    Q   V   V   S   I   S   G   M   G   G   I   G   K   T   T   L   A   R
              CAA GTA GTT TCC ATA TCT GGG ATG GGT GGT ATT GGA AAA ACC ACT CTT GCC AGA
Kranti         Q   V   V   S   I   S   G   M   G   G   I   G   K   T   T   L   A   R
              CAA GTA GTT TCC ATA TCT GGG ATG GGT GGT ATT GGA AAA ACC ACT CTT GCC AGA
Tumida         Q   V   V   S   I   S   G   M   G   G   I   G   K   T   T   L   A   R
              CAA GTA GTT TCC ATA TCT GGG ATG GGT GGT ATT GGA AAA ACC ACT CTT GCC AGA

              600 610 620 630 640
Donskaja-IV    Q   V   F   H   H   D   I   V   R   S   H   F   D   G   F   A   W   I
              CAA GTT TTT CAT CAT GAC ATT GTT CGA AGT CAT TTT GAT GGA TTC GCA TGG ATC
Cutlass        Q   V   F   H   H   D   I   V   R   S   H   F   D   G   F   A   W   I
              CAA GTT TTT CAT CAT GAC ATT GTT CGA AGT CAT TTT GAT GGA TTC GCA TGG ATC
Varuna         Q   V   F   H   H   D   I   V   R   S   H   F   D   G   F   A   W   I
              CAA GTT TTT CAT CAT GAC ATT GTT CGA AGT CAT TTT GAT GGA TTC GCA TGG ATC
Heera          Q   V   F   H   H   D   I   V   R   S   H   F   D   G   F   A   W   I
              CAA GTT TTT CAT CAT GAC ATT GTT CGA AGT CAT TTT GAT GGA TTC GCA TGG ATC
Skorospieka    Q   V   F   H   H   D   I   V   R   S   H   F   D   G   F   A   W   I
              CAA GTT TTT CAT CAT GAC ATT GTT CGA AGT CAT TTT GAT GGA TTC GCA TGG ATC
Kranti         Q   V   F   H   H   D   I   V   R   S   H   F   D   G   F   A   W   I
              CAA GTT TTT CAT CAT GAC ATT GTT CGA AGT CAT TTT GAT GGA TTC GCA TGG ATC
Tumida         Q   V   F   H   H   D   I   V   R   S   H   F   D   G   F   A   W   I
              CAA GTT TTT CAT CAT GAC ATT GTT CGA AGT CAT TTT GAT GGA TTC GCA TGG ATC
```

650 660 670 680 690 700

| | | | | | | | | | | | | | | | | | | |
|---|---|---|---|---|---|---|---|---|---|---|---|---|---|---|---|---|---|---|
| Donskaja-IV | C | V | S | Q | D | F | K | R | K | D | I | W | Q | K | L | L | Q | D |
| | TGC | GTT | TCG | CAA | GAT | TTT | AAG | CGG | AAG | GAT | ATT | TGG | CAA | AAG | TTA | TTG | CAA | GAC |
| Cutlass | C | V | S | Q | D | F | K | R | K | D | I | W | Q | K | L | L | Q | D |
| | TGC | GTT | TCG | CAA | GAT | TTT | AAG | CGG | AAG | GAT | ATT | TGG | CAA | AAG | TTA | TTG | CAA | GAC |
| Varuna | C | V | S | Q | D | C | K | R | M | D | I | W | Q | K | L | L | Q | D |
| | TGC | GTT | TCG | CAA | GAT | TGT | AAG | CGG | ATG | GAT | ATT | TGG | CAA | AAG | TTA | TTG | CAA | GAC |
| Heera | C | V | S | Q | D | C | K | R | M | D | I | W | Q | K | L | L | Q | D |
| | TGC | GTT | TCG | CAA | GAT | TGT | AAG | CGG | ATG | GAT | ATT | TGG | CAA | AAG | TTA | TTG | CAA | GAC |
| Skorospieka | C | V | S | Q | D | C | K | R | M | D | I | W | Q | K | L | L | Q | D |
| | TGC | GTT | TCG | CAA | GAT | TGT | AAG | CGG | ATG | GAT | ATT | TGG | CAA | AAG | TTA | TTG | CAA | GAC |
| Kranti | C | V | S | Q | D | C | K | R | M | D | I | W | Q | K | L | L | Q | D |
| | TGC | GTT | TCG | CAA | GAT | TGT | AAG | CGG | ATG | GAT | ATT | TGG | CAA | AAG | TTA | TTG | CAA | GAC |
| Tumida | C | V | S | Q | D | C | K | R | M | D | I | W | Q | K | L | L | Q | D |
| | TGC | GTT | TCG | CAA | GAT | TGT | AAG | CGG | ATG | GAT | ATT | TGG | CAA | AAG | TTA | TTG | CAA | GAC |

710 720 730 740 750

| | | | | | | | | | | | | | | | | | | |
|---|---|---|---|---|---|---|---|---|---|---|---|---|---|---|---|---|---|---|
| Donskaja-IV | L | R | P | H | D | K | G | N | K | H | M | N | E | E | T | L | Q | A |
| | CTC | AGA | CCA | CAT | GAT | AAG | GGC | AAC | AAA | CAT | ATG | AAT | GAA | GAA | ACA | CTC | CAG | GCT |
| Cutlass | L | R | P | H | D | K | G | N | K | H | M | N | E | E | T | L | Q | A |
| | CTC | AGA | CCA | CAT | GAT | AAG | GGC | AAC | AAA | CAT | ATG | AAT | GAA | GAA | ACA | CTC | CAG | GCT |
| Varuna | L | R | P | Q | D | K | G | I | E | Q | M | N | E | R | T | L | Q | A |
| | CTT | AGA | CCT | CAA | GAT | AAG | GGC | ATC | GAA | CAA | ATG | AAT | GAG | CGT | ACA | CTC | CAG | GCT |
| Heera | L | R | P | Q | D | K | G | I | E | Q | M | N | E | R | T | L | Q | A |
| | CTT | AGA | CCT | CAA | GAT | AAG | GGC | ATC | GAA | CAA | ATG | AAT | GAG | CGT | ACA | CTC | CAG | GCT |
| Skorospieka | L | R | P | Q | D | K | G | I | E | Q | M | N | E | R | T | L | Q | A |
| | CTT | AGA | CCT | CAA | GAT | AAG | GGC | ATC | GAA | CAA | ATG | AAT | GAG | CGT | ACA | CTC | CAG | GCT |
| Kranti | L | R | P | Q | D | K | G | I | E | Q | M | N | E | R | T | L | Q | A |
| | CTT | AGA | CCT | CAA | GAT | AAG | GGC | ATC | GAA | CAA | ATG | AAT | GAG | CGT | ACA | CTC | CAG | GCT |
| Tumida | L | R | P | Q | D | K | G | I | E | Q | M | N | E | R | T | L | Q | A |
| | CTT | AGA | CCT | CAA | GAT | AAG | GGC | ATC | GAA | CAA | ATG | AAT | GAG | CGT | ACA | CTC | CAG | GCT |

760 770 780 790 800 810

| | | | | | | | | | | | | | | | | | | |
|---|---|---|---|---|---|---|---|---|---|---|---|---|---|---|---|---|---|---|
| Donskaja-IV | E | L | F | H | L | L | E | T | S | K | Y | L | I | V | M | D | D | V |
| | GAG | CTC | TTT | CAC | TTG | TTG | GAA | ACA | TCT | AAA | TAT | TTA | ATT | GTC | ATG | GAT | GAT | GTA |
| Cutlass | E | L | F | H | L | L | E | T | S | K | Y | L | I | V | M | D | D | V |
| | GAG | CTC | TTT | CAC | TTG | TTG | GAA | ACA | TCT | AAA | TAT | TTA | ATT | GTC | ATG | GAT | GAT | GTA |
| Varuna | K | I | F | H | L | L | E | T | S | K | Y | L | I | V | M | D | D | V |
| | AAG | ATC | TTT | CAC | TTG | TTG | GAA | ACA | TCT | AAA | TAT | TTA | ATT | GTC | ATG | GAT | GAT | GTA |
| Heera | K | I | F | H | L | L | E | T | S | K | Y | L | I | V | M | D | D | V |
| | AAG | ATC | TTT | CAC | TTG | TTG | GAA | ACA | TCT | AAA | TAT | TTA | ATT | GTC | ATG | GAT | GAT | GTA |
| Skorospieka | K | I | F | H | L | L | E | T | S | K | Y | L | I | V | M | D | D | V |
| | AAG | ATC | TTT | CAC | TTG | TTG | GAA | ACA | TCT | AAA | TAT | TTA | ATT | GTC | ATG | GAT | GAT | GTA |
| Kranti | K | I | F | H | L | L | E | T | S | K | Y | L | I | V | M | D | D | V |
| | AAG | ATC | TTT | CAC | TTG | TTG | GAA | ACA | TCT | AAA | TAT | TTA | ATT | GTC | ATG | GAT | GAT | GTA |
| Tumida | K | I | F | H | L | L | E | T | S | K | Y | L | I | V | M | D | D | V |
| | AAG | ATC | TTT | CAC | TTG | TTG | GAA | ACA | TCT | AAA | TAT | TTA | ATT | GTC | ATG | GAT | GAT | GTA |

820 830 840 850 860

| | | | | | | | | | | | | | | | | | | |
|---|---|---|---|---|---|---|---|---|---|---|---|---|---|---|---|---|---|---|
| Donskaja-IV | W | K | K | E | D | W | D | V | I | K | D | V | F | P | Q | E | R | G |
| | TGG | AAA | AAA | GAA | GAC | TGG | GAT | GTA | ATA | AAA | GAC | GTT | TTC | CCC | CAA | GAA | AGA | GGT |
| Cutlass | W | K | K | E | D | W | D | V | I | K | D | V | F | P | Q | E | R | G |
| | TGG | AAA | AAA | GAA | GAC | TGG | GAT | GTA | ATA | AAA | GAC | GTT | TTC | CCC | CAA | GAA | AGA | GGT |
| Varuna | W | K | K | E | D | W | D | I | I | K | G | V | F | P | Q | E | R | G |
| | TGG | AAA | AAA | GAA | GAC | TGG | GAT | ATA | ATA | AAA | GGC | GTT | TTC | CCC | CAA | GAA | AGA | GGT |
| Heera | W | K | K | E | D | W | D | I | I | K | G | V | F | P | Q | E | R | G |
| | TGG | AAA | AAA | GAA | GAC | TGG | GAT | ATA | ATA | AAA | GGC | GTT | TTC | CCC | CAA | GAA | AGA | GGT |
| Skorospieka | W | K | K | E | D | W | D | I | I | K | G | V | F | P | Q | E | R | G |
| | TGG | AAA | AAA | GAA | GAC | TGG | GAT | ATA | ATA | AAA | GGC | GTT | TTC | CCC | CAA | GAA | AGA | GGT |
| Kranti | W | K | K | E | D | W | D | I | I | K | G | V | F | P | Q | E | R | G |
| | TGG | AAA | AAA | GAA | GAC | TGG | GAT | ATA | ATA | AAA | GGC | GTT | TTC | CCC | CAA | GAA | AGA | GGT |
| Tumida | W | K | K | E | D | W | D | I | I | K | G | V | F | P | Q | E | R | G |
| | TGG | AAA | AAA | GAA | GAC | TGG | GAT | ATA | ATA | AAA | GGC | GTT | TTC | CCC | CAA | GAA | AGA | GGT |

870 880 890 900 910

| | | | | | | | | | | | | | | | | | | |
|---|---|---|---|---|---|---|---|---|---|---|---|---|---|---|---|---|---|---|
| Donskaja-IV | W | K | M | I | L | T | S | R | D | E | G | V | G | L | H | V | N | P |
| | TGG | AAG | ATG | ATT | CTT | ACT | TCA | CGT | GAT | GAA | GGT | GTT | GGT | TTG | CAT | GTA | AAC | CCA |
| Cutlass | W | K | M | I | L | T | S | R | D | E | G | V | G | L | H | V | N | P |
| | TGG | AAG | ATG | ATT | CTT | ACT | TCA | CGT | GAT | GAA | GGT | GTT | GGT | TTG | CAT | GTA | AAC | CCA |
| Varuna | W | K | M | I | L | T | S | R | D | E | S | V | G | F | H | A | D | R |
| | TGG | AAA | ATG | ATT | CTT | ACT | TCA | CGT | GAT | GAA | AGT | GTT | GGT | TTC | CAT | GCA | GAC | CGA |
| Heera | W | K | M | I | L | T | S | R | D | E | S | V | G | F | H | A | D | R |
| | TGG | AAA | ATG | ATT | CTT | ACT | TCA | CGT | GAT | GAA | AGT | GTT | GGT | TTC | CAT | GCA | GAC | CGA |
| Skorospieka | W | K | M | I | L | T | S | R | D | E | S | V | G | F | H | A | D | R |
| | TGG | AAA | ATG | ATT | CTT | ACT | TCA | CGT | GAT | GAA | AGT | GTT | GGT | TTC | CAT | GCA | GAC | CGA |
| Kranti | W | K | M | I | L | T | S | R | D | E | S | V | G | F | H | A | D | R |
| | TGG | AAA | ATG | ATT | CTT | ACT | TCA | CGT | GAT | GAA | AGT | GTT | GGT | TTC | CAT | GCA | GAC | CGA |
| Tumida | W | K | M | I | L | T | S | R | D | E | S | V | G | F | H | A | D | R |
| | TGG | AAA | ATG | ATT | CTT | ACT | TCA | CGT | GAT | GAA | AGT | GTT | GGT | TTC | CAT | GCA | GAC | CGA |

920 930 940 950 960 970

| | | | | | | | | | | | | | | | | | | |
|---|---|---|---|---|---|---|---|---|---|---|---|---|---|---|---|---|---|---|
| Donskaja-IV | K | S | F | L | F | K | P | K | I | L | T | P | E | E | S | W | K | L |
| | AAA | TCT | TTC | CTC | TTT | AAA | CCA | AAA | ATC | CTT | ACT | CCT | GAA | GAA | AGT | TGG | AAG | CTA |
| Cutlass | K | S | F | L | F | K | P | K | I | L | T | P | E | E | S | W | K | L |
| | AAA | TCT | TTC | CTC | TTT | AAA | CCA | AAA | ATC | CTT | ACT | CCT | GAA | GAA | AGT | TGG | AAG | CTA |
| Varuna | T | C | F | P | F | R | P | K | I | L | T | P | E | E | S | W | K | L |
| | ACA | TGT | TTT | CCC | TTT | AGA | CCA | AAA | ATC | CTT | ACT | CCT | GAA | GAA | AGT | TGG | AAG | CTA |
| Heera | T | C | F | P | F | R | P | K | I | L | T | P | E | E | S | W | K | L |
| | ACA | TGT | TTT | CCC | TTT | AGA | CCA | AAA | ATC | CTT | ACT | CCT | GAA | GAA | AGT | TGG | AAG | CTA |
| Skorospieka | T | C | F | P | F | R | P | K | I | L | T | P | E | E | S | W | K | L |
| | ACA | TGT | TTT | CCC | TTT | AGA | CCA | AAA | ATC | CTT | ACT | CCT | GAA | GAA | AGT | TGG | AAG | CTA |
| Kranti | T | C | F | P | F | R | P | K | I | L | T | P | E | E | S | W | K | L |
| | ACA | TGT | TTT | CCC | TTT | AGA | CCA | AAA | ATC | CTT | ACT | CCT | GAA | GAA | AGT | TGG | AAG | CTA |
| Tumida | T | C | F | P | F | R | P | K | I | L | T | P | E | E | S | W | K | L |
| | ACA | TGT | TTT | CCC | TTT | AGA | CCA | AAA | ATC | CTT | ACT | CCT | GAA | GAA | AGT | TGG | AAG | CTA |

980 990 1000 1010 1020

| | | | | | | | | | | | | | | | | | | |
|---|---|---|---|---|---|---|---|---|---|---|---|---|---|---|---|---|---|---|
| Donskaja-IV | C | E | S | I | V | F | S | R | R | E | I | T | E | F | I | V | D | E |
| | TGT | GAG | AGC | ATA | GTA | TTC | TCT | AGA | AGA | GAG | ATA | ACC | GAA | TTT | ATT | GTT | GAT | GAA |
| Cutlass | C | E | S | I | V | F | S | R | R | E | I | T | E | F | I | V | D | E |
| | TGT | GAG | AGC | ATA | GTA | TTC | TCT | AGA | AGA | GAG | ATA | ACC | GAA | TTT | ATT | GTT | GAT | GAA |
| Varuna | C | E | G | I | V | F | P | R | R | E | I | T | E | F | N | V | D | E |
| | TGT | GAG | GGC | ATA | GTA | TTC | CCT | AGA | AGA | GAG | ATA | ACG | GAA | TTT | AAT | GTT | GAT | GAA |
| Heera | C | E | G | I | V | F | P | R | R | E | I | T | E | F | N | V | D | E |
| | TGT | GAG | GGC | ATA | GTA | TTC | CCT | AGA | AGA | GAG | ATA | ACG | GAA | TTT | AAT | GTT | GAT | GAA |
| Skorospieka | C | E | G | I | V | F | P | R | R | E | I | T | E | F | N | V | D | E |
| | TGT | GAG | GGC | ATA | GTA | TTC | CCT | AGA | AGA | GAG | ATA | ACG | GAA | TTT | AAT | GTT | GAT | GAA |
| Kranti | C | E | G | I | V | F | P | R | R | E | I | T | E | F | N | V | D | E |
| | TGT | GAG | GGC | ATA | GTA | TTC | CCT | AGA | AGA | GAG | ATA | ACG | GAA | TTT | AAT | GTT | GAT | GAA |
| Tumida | C | E | G | I | V | F | P | R | R | E | I | T | E | F | N | V | D | E |
| | TGT | GAG | GGC | ATA | GTA | TTC | CCT | AGA | AGA | GAG | ATA | ACG | GAA | TTT | AAT | GTT | GAT | GAA |

1030 1040 1050 1060 1070 1080

| | | | | | | | | | | | | | | | | | | |
|---|---|---|---|---|---|---|---|---|---|---|---|---|---|---|---|---|---|---|
| Donskaja-IV | E | L | E | A | M | G | R | K | M | V | K | Y | C | G | G | L | P | L |
| | GAA | CTA | GAA | GCC | ATG | GGA | AGG | AAA | ATG | GTC | AAA | TAT | TGT | GGA | GGA | CTA | CCA | TTA |
| Cutlass | E | L | E | A | M | G | R | K | M | V | K | Y | C | G | G | L | P | L |
| | GAA | CTA | GAA | GCC | ATG | GGA | AGG | AAA | ATG | GTC | AAA | TAT | TGT | GGA | GGA | CTA | CCA | TTA |
| Varuna | E | L | E | A | M | G | K | K | M | V | K | Y | C | G | G | L | P | L |
| | GAA | CTA | GAA | GCC | ATG | GGT | AAG | AAA | ATG | GTC | AAA | TAT | TGT | GGA | GGA | CTA | CCA | TTA |
| Heera | E | L | E | A | M | G | K | K | M | V | K | Y | C | G | G | L | P | L |
| | GAA | CTA | GAA | GCC | ATG | GGT | AAG | AAA | ATG | GTC | AAA | TAT | TGT | GGA | GGA | CTA | CCA | TTA |
| Skorospieka | E | L | E | A | M | G | K | K | M | V | K | Y | C | G | G | L | P | L |
| | GAA | CTA | GAA | GCC | ATG | GGT | AAG | AAA | ATG | GTC | AAA | TAT | TGT | GGA | GGA | CTA | CCA | TTA |
| Kranti | E | L | E | A | M | G | K | K | M | V | K | Y | C | G | G | L | P | L |
| | GAA | CTA | GAA | GCC | ATG | GGT | AAG | AAA | ATG | GTC | AAA | TAT | TGT | GGA | GGA | CTA | CCA | TTA |
| Tumida | E | L | E | A | M | G | K | K | M | V | K | Y | C | G | G | L | P | L |
| | GAA | CTA | GAA | GCC | ATG | GGT | AAG | AAA | ATG | GTC | AAA | TAT | TGT | GGA | GGA | CTA | CCA | TTA |

1090 1100 1110 1120 1130

| | | | | | | | | | | | | | | | | | | |
|---|---|---|---|---|---|---|---|---|---|---|---|---|---|---|---|---|---|---|
| Donskaja-IV | A | V | K | V | L | G | G | L | L | A | N | K | I | M | V | E | E | W |
| | GCT | GTT | AAA | GTG | TTG | GGA | GGT | TTG | TTA | GCT | AAT | AAA | ATC | ATG | GTT | GAA | GAG | TGG |
| Cutlass | A | V | K | V | L | G | G | L | L | A | N | K | I | M | V | E | E | W |
| | GCT | GTT | AAA | GTG | TTG | GGA | GGT | TTG | TTA | GCT | AAT | AAA | ATC | ATG | GTT | GAA | GAG | TGG |
| Varuna | A | V | K | V | L | G | G | L | L | A | N | K | T | M | V | E | E | W |
| | GCC | GTT | AAA | GTG | CTG | GGA | GGC | TTG | TTA | GCT | AAT | AAA | ACC | ATG | GTT | GAA | GAG | TGG |
| Heera | A | V | K | V | L | G | G | L | L | A | N | K | T | M | V | E | E | W |
| | GCC | GTT | AAA | GTG | CTG | GGA | GGC | TTG | TTA | GCT | AAT | AAA | ACC | ATG | GTT | GAA | GAG | TGG |
| Skorospieka | A | V | K | V | L | G | G | L | L | A | N | K | T | M | V | E | E | W |
| | GCC | GTT | AAA | GTG | CTG | GGA | GGC | TTG | TTA | GCT | AAT | AAA | ACC | ATG | GTT | GAA | GAG | TGG |
| Kranti | A | V | K | V | L | G | G | L | L | A | N | K | T | M | V | E | E | W |
| | GCC | GTT | AAA | GTG | CTG | GGA | GGC | TTG | TTA | GCT | AAT | AAA | ACC | ATG | GTT | GAA | GAG | TGG |
| Tumida | A | V | K | V | L | G | G | L | L | A | N | K | T | M | V | E | E | W |
| | GCC | GTT | AAA | GTG | CTG | GGA | GGC | TTG | TTA | GCT | AAT | AAA | ACC | ATG | GTT | GAA | GAG | TGG |

1140 1150 1160 1170 1180

| | | | | | | | | | | | | | | | | | | |
|---|---|---|---|---|---|---|---|---|---|---|---|---|---|---|---|---|---|---|
| Donskaja-IV | K | R | V | D | D | N | I | K | T | Q | I | V | R | R | D | D | K | N |
| | AAA | AGA | GTA | GAT | GAC | AAT | ATT | AAA | ACT | CAG | ATA | GTT | CGC | AGA | GAT | GAC | AAA | AAT |
| Cutlass | K | R | V | D | D | N | I | K | T | Q | I | V | R | R | D | D | K | N |
| | AAA | AGA | GTA | GAT | GAC | AAT | ATT | AAA | ACT | CAG | ATA | GTT | CGC | AGA | GAT | GAC | AAA | AAT |
| Varuna | K | R | V | D | D | N | I | Q | T | Q | I | V | R | I | D | D | K | N |
| | AAA | AGA | GTA | GAT | GAC | AAT | ATT | CAA | ACT | CAG | ATA | GTT | CGC | ATA | GAT | GAC | AAA | AAT |
| Heera | K | R | V | D | D | N | I | Q | T | Q | I | V | R | I | D | D | K | N |
| | AAA | AGA | GTA | GAT | GAC | AAT | ATT | CAA | ACT | CAG | ATA | GTT | CGC | ATA | GAT | GAC | AAA | AAT |
| Skorospieka | K | R | V | D | D | N | I | Q | T | Q | I | V | R | I | D | D | K | N |
| | AAA | AGA | GTA | GAT | GAC | AAT | ATT | CAA | ACT | CAG | ATA | GTT | CGC | ATA | GAT | GAC | AAA | AAT |
| Kranti | K | R | V | D | D | N | I | Q | T | Q | I | V | R | I | D | D | K | N |
| | AAA | AGA | GTA | GAT | GAC | AAT | ATT | CAA | ACT | CAG | ATA | GTT | CGC | ATA | GAT | GAC | AAA | AAT |
| Tumida | K | R | V | D | D | N | I | Q | T | Q | I | V | R | I | D | D | K | N |
| | AAA | AGA | GTA | GAT | GAC | AAT | ATT | CAA | ACT | CAG | ATA | GTT | CGC | ATA | GAT | GAC | AAA | AAT |

1190 1200 1210 1220 1230 1240

| | | | | | | | | | | | | | | | | | | |
|---|---|---|---|---|---|---|---|---|---|---|---|---|---|---|---|---|---|---|
| Donskaja-IV | Q | D | S | V | Y | R | V | L | S | M | S | Y | E | D | L | P | M | Q |
| | CAA | GAT | TCG | GTT | TAC | CGA | GTA | TTG | TCT | ATG | AGC | TAT | GAA | GAT | TTG | CCA | ATG | CAG |
| Cutlass | Q | D | S | V | Y | R | V | L | S | M | S | Y | E | D | L | P | M | Q |
| | CAA | GAT | TCG | GTT | TAC | CGA | GTA | TTG | TCT | ATG | AGC | TAT | GAA | GAT | TTG | CCA | ATG | CAG |
| Varuna | Q | D | S | V | Y | R | V | L | S | M | S | Y | E | D | L | P | M | Q |
| | CAA | GAT | TCT | GTT | TAC | CGA | GTA | TTG | TCT | ATG | AGC | TAT | GAA | GAT | TTG | CCA | ATG | CAG |
| Heera | Q | D | S | V | Y | R | V | L | S | M | S | Y | E | D | L | P | M | Q |
| | CAA | GAT | TCT | GTT | TAC | CGA | GTA | TTG | TCT | ATG | AGC | TAT | GAA | GAT | TTG | CCA | ATG | CAG |
| Skorospieka | Q | D | S | V | Y | R | V | L | S | M | S | Y | E | D | L | P | M | Q |
| | CAA | GAT | TCT | GTT | TAC | CGA | GTA | TTG | TCT | ATG | AGC | TAT | GAA | GAT | TTG | CCA | ATG | CAG |
| Kranti | Q | D | S | V | Y | R | V | L | S | M | S | Y | E | D | L | P | M | Q |
| | CAA | GAT | TCT | GTT | TAC | CGA | GTA | TTG | TCT | ATG | AGC | TAT | GAA | GAT | TTG | CCA | ATG | CAG |
| Tumida | Q | D | S | V | Y | R | V | L | S | M | S | Y | E | D | L | P | M | Q |
| | CAA | GAT | TCT | GTT | TAC | CGA | GTA | TTG | TCT | ATG | AGC | TAT | GAA | GAT | TTG | CCA | ATG | CAG |

1250 1260 1270 1280 1290

| | | | | | | | | | | | | | | | | | | |
|---|---|---|---|---|---|---|---|---|---|---|---|---|---|---|---|---|---|---|
| Donskaja-IV | L | K | N | C | F | L | Y | L | A | H | F | P | E | D | Y | K | I | E |
| | TTG | AAG | AAT | TGC | TTC | CTC | TAC | CTA | GCT | CAT | TTT | CCA | GAA | GAT | TAC | AAG | ATA | GAA |
| Cutlass | L | K | N | C | F | L | Y | L | A | H | F | P | E | D | Y | K | I | E |
| | TTG | AAG | AAT | TGC | TTC | CTC | TAC | CTA | GCT | CAT | TTT | CCA | GAA | GAT | TAC | AAG | ATA | GAA |
| Varuna | L | K | H | C | F | L | Y | L | A | H | F | P | E | D | Y | E | I | Q |
| | TTA | AAG | CAT | TGC | TTT | CTC | TAC | CTA | GCT | CAT | TTT | CCC | GAA | GAT | TAC | GAG | ATA | CAA |
| Heera | L | K | H | C | F | L | Y | L | A | H | F | P | E | D | Y | E | I | Q |
| | TTA | AAG | CAT | TGC | TTT | CTC | TAC | CTA | GCT | CAT | TTT | CCC | GAA | GAT | TAC | GAG | ATA | CAA |
| Skorospieka | L | K | H | C | F | L | Y | L | A | H | F | P | E | D | Y | E | I | Q |
| | TTA | AAG | CAT | TGC | TTT | CTC | TAC | CTA | GCT | CAT | TTT | CCC | GAA | GAT | TAC | GAG | ATA | CAA |
| Kranti | L | K | H | C | F | L | Y | L | A | H | F | P | E | D | Y | E | I | Q |
| | TTA | AAG | CAT | TGC | TTT | CTC | TAC | CTA | GCT | CAT | TTT | CCC | GAA | GAT | TAC | GAG | ATA | CAA |
| Tumida | L | K | H | C | F | L | Y | L | A | H | F | P | E | D | Y | E | I | Q |
| | TTA | AAG | CAT | TGC | TTT | CTC | TAC | CTA | GCT | CAT | TTT | CCC | GAA | GAT | TAC | GAG | ATA | CAA |

1300 1310 1320 1330 1340 1350

| | | | | | | | | | | | | | | | | | | |
|---|---|---|---|---|---|---|---|---|---|---|---|---|---|---|---|---|---|---|
| Donskaja-IV | A | E | Q | L | Y | Y | Y | W | E | A | E | G | I | I | T | S | S | V |
| | GCA | GAG | CAA | TTG | TAT | TAT | TAC | TGG | GAA | GCA | GAA | GGA | ATA | ATA | ACG | TCA | AGT | GTC |
| Cutlass | A | E | Q | L | Y | Y | Y | W | E | A | E | G | I | I | T | S | S | V |
| | GCA | GAG | CAA | TTG | TAT | TAT | TAC | TGG | GAA | GCA | GAA | GGA | ATA | ATA | ACG | TCA | AGT | GTC |
| Varuna | V | E | R | L | Y | Y | L | W | E | A | E | G | I | I | T | S | S | G |
| | GTG | GAG | AGA | TTG | TAT | TAT | CTC | TGG | GAA | GCA | GAA | GGA | ATA | ATA | ACG | TCA | AGT | GGC |
| Heera | V | E | R | L | Y | Y | L | W | E | A | E | G | I | I | T | S | S | G |
| | GTG | GAG | AGA | TTG | TAT | TAT | CTC | TGG | GAA | GCA | GAA | GGA | ATA | ATA | ACG | TCA | AGT | GGC |
| Skorospieka | V | E | R | L | Y | Y | L | W | E | A | E | G | I | I | T | S | S | G |
| | GTG | GAG | AGA | TTG | TAT | TAT | CTC | TGG | GAA | GCA | GAA | GGA | ATA | ATA | ACG | TCA | AGT | GGC |
| Kranti | V | E | R | L | Y | Y | L | W | E | A | E | G | I | I | T | S | S | G |
| | GTG | GAG | AGA | TTG | TAT | TAT | CTC | TGG | GAA | GCA | GAA | GGA | ATA | ATA | ACG | TCA | AGT | GGC |
| Tumida | V | E | R | L | Y | Y | L | W | E | A | E | G | I | I | T | S | S | G |
| | GTG | GAG | AGA | TTG | TAT | TAT | CTC | TGG | GAA | GCA | GAA | GGA | ATA | ATA | ACG | TCA | AGT | GGC |

1360 1370 1380 1390 1400

| | | | | | | | | | | | | | | | | | | |
|---|---|---|---|---|---|---|---|---|---|---|---|---|---|---|---|---|---|---|
| Donskaja-IV | D | G | E | T | T | R | K | I | G | E | D | Y | I | D | E | L | V | R |
| | GAT | GGA | GAA | ACC | ACT | AGG | AAA | ATT | GGA | GAA | GAC | TAC | ATA | GAC | GAG | CTA | GTA | AGG |
| Cutlass | D | G | E | T | T | R | K | I | G | E | D | Y | I | D | E | L | V | R |
| | GAT | GGA | GAA | ACC | ACT | AGG | AAA | ATT | GGA | GAA | GAC | TAC | ATA | GAC | GAG | CTA | GTA | AGG |
| Varuna | D | G | E | T | T | R | K | I | G | E | E | Y | I | D | E | L | V | R |
| | GAT | GGA | GAA | ACC | ACT | AGG | AAA | ATT | GGA | GAA | GAA | TAC | ATA | GAC | GAG | CTA | GTA | AGG |
| Heera | D | G | E | T | T | R | K | I | G | E | E | Y | I | D | E | L | V | R |
| | GAT | GGA | GAA | ACC | ACT | AGG | AAA | ATT | GGA | GAA | GAA | TAC | ATA | GAC | GAG | CTA | GTA | AGG |
| Skorospieka | D | G | E | T | T | R | K | I | G | E | E | Y | I | D | E | L | V | R |
| | GAT | GGA | GAA | ACC | ACT | AGG | AAA | ATT | GGA | GAA | GAA | TAC | ATA | GAC | GAG | CTA | GTA | AGG |
| Kranti | D | G | E | T | T | R | K | I | G | E | E | Y | I | D | E | L | V | R |
| | GAT | GGA | GAA | ACC | ACT | AGG | AAA | ATT | GGA | GAA | GAA | TAC | ATA | GAC | GAG | CTA | GTA | AGG |
| Tumida | D | G | E | T | T | R | K | I | G | E | E | Y | I | D | E | L | V | R |
| | GAT | GGA | GAA | ACC | ACT | AGG | AAA | ATT | GGA | GAA | GAA | TAC | ATA | GAC | GAG | CTA | GTA | AGG |

1410 1420 1430 1440 1450

| | | | | | | | | | | | | | | | | | | |
|---|---|---|---|---|---|---|---|---|---|---|---|---|---|---|---|---|---|---|
| Donskaja-IV | R | N | M | I | I | G | V | K | E | D | L | S | C | R | W | E | Y | C |
| | AGA | AAT | ATG | ATT | ATT | GGA | GTA | AAA | GAG | GAT | TTG | AGT | TGC | AGA | TGG | GAG | TAT | TGT |
| Cutlass | R | N | M | I | I | G | V | K | E | D | L | S | C | R | W | E | Y | C |
| | AGA | AAT | ATG | ATT | ATT | GGA | GTA | AAA | GAG | GAT | TTG | AGT | TGC | AGA | TGG | GAG | TAT | TGT |
| Varuna | R | N | M | V | I | G | V | K | E | D | L | S | C | K | W | E | Y | C |
| | AGA | AAT | ATG | GTT | ATT | GGA | GTA | AAA | GAG | GAT | TTG | AGT | TGC | AAA | TGG | GAG | TAT | TGT |
| Heera | R | N | M | V | I | G | V | K | E | D | L | S | C | K | W | E | Y | C |
| | AGA | AAT | ATG | GTT | ATT | GGA | GTA | AAA | GAG | GAT | TTG | AGT | TGC | AAA | TGG | GAG | TAT | TGT |
| Skorospieka | R | N | M | V | I | G | V | K | E | D | L | S | C | K | W | E | Y | C |
| | AGA | AAT | ATG | GTT | ATT | GGA | GTA | AAA | GAG | GAT | TTG | AGT | TGC | AAA | TGG | GAG | TAT | TGT |
| Kranti | R | N | M | V | I | G | V | K | E | D | L | S | C | K | W | E | Y | C |
| | AGA | AAT | ATG | GTT | ATT | GGA | GTA | AAA | GAG | GAT | TTG | AGT | TGC | AAA | TGG | GAG | TAT | TGT |
| Tumida | R | N | M | V | I | G | V | K | E | D | L | S | C | K | W | E | Y | C |
| | AGA | AAT | ATG | GTT | ATT | GGA | GTA | AAA | GAG | GAT | TTG | AGT | TGC | AAA | TGG | GAG | TAT | TGT |

1460 1470 1480 1490 1500 1510

| | | | | | | | | | | | | | | | | | | |
|---|---|---|---|---|---|---|---|---|---|---|---|---|---|---|---|---|---|---|
| Donskaja-IV | Q | M | H | D | M | M | R | E | V | C | L | S | K | A | K | E | E | N |
| | CAA | ATG | CAT | GAC | ATG | ATG | AGA | GAA | GTA | TGT | TTA | TCT | AAA | GCC | AAA | GAA | GAG | AAC |
| Cutlass | Q | M | H | D | M | M | R | E | V | C | L | S | K | A | K | E | E | N |
| | CAA | ATG | CAT | GAC | ATG | ATG | AGA | GAA | GTA | TGT | TTA | TCT | AAA | GCC | AAA | GAA | GAG | AAC |
| Varuna | Q | M | H | D | M | M | R | E | V | C | L | Y | K | A | K | E | E | N |
| | CAA | ATG | CAT | GAC | ATG | ATG | AGA | GAA | GTA | TGT | TTA | TAT | AAA | GCC | AAA | GAA | GAG | AAC |
| Heera | Q | M | H | D | M | M | R | E | V | C | L | Y | K | A | K | E | E | N |
| | CAA | ATG | CAT | GAC | ATG | ATG | AGA | GAA | GTA | TGT | TTA | TAT | AAA | GCC | AAA | GAA | GAG | AAC |
| Skorospieka | Q | M | H | D | M | M | R | E | V | C | L | Y | K | A | K | E | E | N |
| | CAA | ATG | CAT | GAC | ATG | ATG | AGA | GAA | GTA | TGT | TTA | TAT | AAA | GCC | AAA | GAA | GAG | AAC |
| Kranti | Q | M | H | D | M | M | R | E | V | C | L | Y | K | A | K | E | E | N |
| | CAA | ATG | CAT | GAC | ATG | ATG | AGA | GAA | GTA | TGT | TTA | TAT | AAA | GCC | AAA | GAA | GAG | AAC |
| Tumida | Q | M | H | D | M | M | R | E | V | C | L | Y | K | A | K | E | E | N |
| | CAA | ATG | CAT | GAC | ATG | ATG | AGA | GAA | GTA | TGT | TTA | TAT | AAA | GCC | AAA | GAA | GAG | AAC |

1520 1530 1540 1550 1560

| | | | | | | | | | | | | | | | | | | |
|---|---|---|---|---|---|---|---|---|---|---|---|---|---|---|---|---|---|---|
| Donskaja-IV | F | L | Q | F | I | K | V | P | T | T | S | T | S | T | I | N | A | H |
| | TTT | CTT | CAG | TTT | ATC | AAA | GTC | CCT | ACT | ACT | TCC | ACC | TCC | ACC | ATC | AAT | GCT | CAT |
| Cutlass | F | L | Q | F | I | K | V | P | T | T | S | T | S | T | I | N | A | H |
| | TTT | CTT | CAG | TTT | ATC | AAA | GTC | CCT | ACT | ACT | TCC | ACC | TCC | ACC | ATC | AAT | GCT | CAT |
| Varuna | F | L | L | I | I | K | V | P | T | S | S | T | S | T | I | N | A | Q |
| | TTT | CTT | CTG | ATT | ATC | AAA | GTC | CCT | ACT | TCT | TCC | ACC | TCC | ACC | ATC | AAT | GCT | CAA |
| Heera | F | L | L | I | I | K | V | P | T | S | S | T | S | T | I | N | A | Q |
| | TTT | CTT | CTG | ATT | ATC | AAA | GTC | CCT | ACT | TCT | TCC | ACC | TCC | ACC | ATC | AAT | GCT | CAA |
| Skorospieka | F | L | L | I | I | K | V | P | T | S | S | T | S | T | I | N | A | Q |
| | TTT | CTT | CTG | ATT | ATC | AAA | GTC | CCT | ACT | TCT | TCC | ACC | TCC | ACC | ATC | AAT | GCT | CAA |
| Kranti | F | L | L | I | I | K | V | P | T | S | S | T | S | T | I | N | A | Q |
| | TTT | CTT | CTG | ATT | ATC | AAA | GTC | CCT | ACT | TCT | TCC | ACC | TCC | ACC | ATC | AAT | GCT | CAA |
| Tumida | F | L | L | I | I | K | V | P | T | S | S | T | S | T | I | N | A | Q |
| | TTT | CTT | CTG | ATT | ATC | AAA | GTC | CCT | ACT | TCT | TCC | ACC | TCC | ACC | ATC | AAT | GCT | CAA |

1570 1580 1590 1600 1610 1620

| | | | | | | | | | | | | | | | | | | |
|---|---|---|---|---|---|---|---|---|---|---|---|---|---|---|---|---|---|---|
| Donskaja-IV | T | P | T | R | S | R | R | L | V | V | H | G | G | G | N | A | F | D |
| | ACT | CCT | ACC | AGA | TCT | CGC | AGA | CTC | GTA | GTA | CAC | GGT | GGT | GGT | AAT | GCA | TTT | GAT |
| Cutlass | T | P | T | R | S | R | R | L | V | V | H | G | G | G | N | A | F | D |
| | ACT | CCT | ACC | AGA | TCT | CGC | AGA | CTC | GTA | GTA | CAC | GGT | GGT | GGT | AAT | GCA | TTT | GAT |
| Varuna | S | H | T | G | S | R | R | L | V | V | H | G | G | G | N | A | F | D |
| | AGT | CAT | ACG | GGA | TCT | CGC | AGA | CTC | GTA | GTA | CAC | GGT | GGT | GGT | AAT | GCA | TTT | GAT |
| Heera | S | H | T | G | S | R | R | L | V | V | H | G | G | G | N | A | F | D |
| | AGT | CAT | ACG | GGA | TCT | CGC | AGA | CTC | GTA | GTA | CAC | GGT | GGT | GGT | AAT | GCA | TTT | GAT |
| Skorospieka | S | H | T | G | S | R | R | L | V | V | H | G | G | G | N | A | F | D |
| | AGT | CAT | ACG | GGA | TCT | CGC | AGA | CTC | GTA | GTA | CAC | GGT | GGT | GGT | AAT | GCA | TTT | GAT |
| Kranti | S | H | T | G | S | R | R | L | V | V | H | G | G | G | N | A | F | D |
| | AGT | CAT | ACG | GGA | TCT | CGC | AGA | CTC | GTA | GTA | CAC | GGT | GGT | GGT | AAT | GCA | TTT | GAT |
| Tumida | S | H | T | G | S | R | R | L | V | V | H | G | G | G | N | A | F | D |
| | AGT | CAT | ACG | GGA | TCT | CGC | AGA | CTC | GTA | GTA | CAC | GGT | GGT | GGT | AAT | GCA | TTT | GAT |

1630 1640 1650 1660 1670

| | | | | | | | | | | | | | | | | | | |
|---|---|---|---|---|---|---|---|---|---|---|---|---|---|---|---|---|---|---|
| Donskaja-IV | M | L | E | R | K | N | N | Q | K | A | R | S | V | L | G | F | G | L |
| | ATG | CTG | GAA | CGT | AAA | AAC | AAT | CAA | AAA | GCT | AGA | TCT | GTT | TTG | GGT | TTT | GGA | CTC |
| Cutlass | M | L | E | R | K | N | N | Q | K | A | R | S | V | L | G | F | G | L |
| | ATG | CTG | GAA | CGT | AAA | AAC | AAT | CAA | AAA | GCT | AGA | TCT | GTT | TTG | GGT | TTT | GGA | CTC |
| Varuna | M | L | E | R | K | N | N | Q | K | A | R | S | V | L | G | L | G | L |
| | ATG | CTG | GAA | CGT | AAA | AAC | AAT | CAA | AAA | GCT | AGA | TCT | GTT | TTG | GGG | TTA | GGA | CTC |
| Heera | M | L | E | R | K | N | N | Q | K | A | R | S | V | L | G | L | G | L |
| | ATG | CTG | GAA | CGT | AAA | AAC | AAT | CAA | AAA | GCT | AGA | TCT | GTT | TTG | GGG | TTA | GGA | CTC |
| Skorospieka | M | L | E | R | K | N | N | Q | K | A | R | S | V | L | G | L | G | L |
| | ATG | CTG | GAA | CGT | AAA | AAC | AAT | CAA | AAA | GCT | AGA | TCT | GTT | TTG | GGG | TTA | GGA | CTC |
| Kranti | M | L | E | R | K | N | N | Q | K | A | R | S | V | L | G | L | G | L |
| | ATG | CTG | GAA | CGT | AAA | AAC | AAT | CAA | AAA | GCT | AGA | TCT | GTT | TTG | GGG | TTA | GGA | CTC |
| Tumida | M | L | E | R | K | N | N | Q | K | A | R | S | V | L | G | L | G | L |
| | ATG | CTG | GAA | CGT | AAA | AAC | AAT | CAA | AAA | GCT | AGA | TCT | GTT | TTG | GGG | TTA | GGA | CTC |

1680 1690 1700 1710 1720

| | | | | | | | | | | | | | | | | | | |
|---|---|---|---|---|---|---|---|---|---|---|---|---|---|---|---|---|---|---|
| Donskaja-IV | D | S | N | L | W | K | Q | S | G | Q | G | F | Q | N | L | L | L | L |
| | GAC | AGC | AAC | TTG | TGG | AAG | CAG | TCA | GGT | CAA | GGA | TTC | CAA | AAT | TTA | CTG | TTA | CTT |
| Cutlass | D | S | N | L | W | K | Q | S | G | Q | G | F | Q | N | L | L | L | L |
| | GAC | AGC | AAC | TTG | TGG | AAG | CAG | TCA | GGT | CAA | GGA | TTC | CAA | AAT | TTA | CTG | TTA | CTT |
| Varuna | D | S | N | F | W | K | Q | S | G | Q | G | F | Q | N | L | L | L | L |
| | GAC | AGC | AAC | TTT | TGG | AAG | CAG | TCA | GGT | CAA | GGA | TTC | CAA | AAT | TTA | CTG | TTA | CTT |
| Heera | D | S | N | F | W | K | Q | S | G | Q | G | F | Q | N | L | L | L | L |
| | GAC | AGC | AAC | TTT | TGG | AAG | CAG | TCA | GGT | CAA | GGA | TTC | CAA | AAT | TTA | CTG | TTA | CTT |
| Skorospieka | D | S | N | F | W | K | Q | S | G | Q | G | F | Q | N | L | L | L | L |
| | GAC | AGC | AAC | TTT | TGG | AAG | CAG | TCA | GGT | CAA | GGA | TTC | CAA | AAT | TTA | CTG | TTA | CTT |
| Kranti | D | S | N | F | W | K | Q | S | G | Q | G | F | Q | N | L | L | L | L |
| | GAC | AGC | AAC | TTT | TGG | AAG | CAG | TCA | GGT | CAA | GGA | TTC | CAA | AAT | TTA | CTG | TTA | CTT |
| Tumida | D | S | N | F | W | K | Q | S | G | Q | G | F | Q | N | L | L | L | L |
| | GAC | AGC | AAC | TTT | TGG | AAG | CAG | TCA | GGT | CAA | GGA | TTC | CAA | AAT | TTA | CTG | TTA | CTT |

1730 1740 1750 1760 1770 1780

| | | | | | | | | | | | | | | | | | | |
|---|---|---|---|---|---|---|---|---|---|---|---|---|---|---|---|---|---|---|
| Donskaja-IV | R | V | L | D | L | S | L | D | Y | K | I | D | S | K | G | W | R | I |
| | AGG | GTC | TTA | GAT | TTG | AGT | TTA | GAT | TAT | AAA | ATA | GAC | TCT | AAA | GGG | TGG | AGA | ATA |
| Cutlass | R | V | L | D | L | S | L | D | Y | K | I | D | S | K | G | W | R | I |
| | AGG | GTC | TTA | GAT | TTG | AGT | TTA | GAT | TAT | AAA | ATA | GAC | TCT | AAA | GGG | TGG | AGA | ATA |
| Varuna | R | V | L | D | L | S | L | D | Y | K | I | D | S | K | G | W | R | I |
| | AGG | GTC | TTA | GAT | TTG | AGT | TTA | GAT | TAT | AAA | ATA | GAC | TCT | AAA | GGG | TGG | AGA | ATA |
| Heera | R | V | L | D | L | S | L | D | Y | K | I | D | S | K | G | W | R | I |
| | AGG | GTC | TTA | GAT | TTG | AGT | TTA | GAT | TAT | AAA | ATA | GAC | TCT | AAA | GGG | TGG | AGA | ATA |
| Skorospieka | R | V | L | D | L | S | L | D | Y | K | I | D | S | K | G | W | R | I |
| | AGG | GTC | TTA | GAT | TTG | AGT | TTA | GAT | TAT | AAA | ATA | GAC | TCT | AAA | GGG | TGG | AGA | ATA |
| Kranti | R | V | L | D | L | S | L | D | Y | K | I | D | S | K | G | W | R | I |
| | AGG | GTC | TTA | GAT | TTG | AGT | TTA | GAT | TAT | AAA | ATA | GAC | TCT | AAA | GGG | TGG | AGA | ATA |
| Tumida | R | V | L | D | L | S | L | D | Y | K | I | D | S | K | G | W | R | I |
| | AGG | GTC | TTA | GAT | TTG | AGT | TTA | GAT | TAT | AAA | ATA | GAC | TCT | AAA | GGG | TGG | AGA | ATA |

1790 1800 1810 1820 1830

| | | | | | | | | | | | | | | | | | | |
|---|---|---|---|---|---|---|---|---|---|---|---|---|---|---|---|---|---|---|
| Donskaja-IV | P | S | S | I | G | K | L | I | H | L | R | Y | L | R | L | E | M | G |
| | CCT | TCG | AGC | ATT | GGA | AAA | CTC | ATT | CAT | TTG | AGA | TAT | TTG | AGA | TTA | GAG | ATG | GGT |
| Cutlass | P | S | S | I | - | - | - | - | H | L | R | Y | L | R | L | E | M | G |
| | CCT | TCG | AGC | ATT | --- | --- | --- | --- | CAT | TTG | AGA | TAT | TTG | AGA | TTA | GAG | ATG | GGT |
| Varuna | P | S | S | I | - | - | - | - | H | L | R | Y | L | R | L | Y | M | G |
| | CCT | TCG | AGC | ATT | --- | --- | --- | --- | CAT | TTG | AGA | TAT | TTG | AGA | TTA | TAT | ATG | GGT |
| Heera | P | S | S | I | - | - | - | - | H | L | R | Y | L | R | L | Y | M | G |
| | CCT | TCG | AGC | ATT | --- | --- | --- | --- | CAT | TTG | AGA | TAT | TTG | AGA | TTA | TAT | ATG | GGT |
| Skorospieka | P | S | S | I | - | - | - | - | H | L | R | Y | L | R | L | Y | M | G |
| | CCT | TCG | AGC | ATT | --- | --- | --- | --- | CAT | TTG | AGA | TAT | TTG | AGA | TTA | TAT | ATG | GGT |
| Kranti | P | S | S | I | - | - | - | - | H | L | R | Y | L | R | L | Y | M | G |
| | CCT | TCG | AGC | ATT | --- | --- | --- | --- | CAT | TTG | AGA | TAT | TTG | AGA | TTA | TAT | ATG | GGT |
| Tumida | P | S | S | I | - | - | - | - | H | L | R | Y | L | R | L | Y | M | G |
| | CCT | TCG | AGC | ATT | --- | --- | --- | --- | CAT | TTG | AGA | TAT | TTG | AGA | TTA | TAT | ATG | GGT |

1840 1850 1860 1870 1880 1890

| | | | | | | | | | | | | | | | | | | |
|---|---|---|---|---|---|---|---|---|---|---|---|---|---|---|---|---|---|---|
| Donskaja-IV | H | A | T | H | V | P | S | S | L | R | N | L | K | L | L | I | Y | L |
| | CAT | GCA | ACT | CAT | GTA | CCT | TCT | TCT | TTA | CGG | AAT | CTA | AAG | CTT | CTA | ATC | TAT | TTG |
| Cutlass | H | A | T | H | V | P | S | S | L | R | N | L | K | L | L | I | Y | L |
| | CAT | GCA | ACT | CAT | GTA | CCT | TCT | TCT | TTA | CGG | AAT | CTA | AAG | CTT | CTA | ATC | TAT | TTG |
| Varuna | H | A | T | H | V | P | S | S | L | R | N | L | K | L | L | I | Y | L |
| | CAT | GCA | ACT | CAT | GTA | CCT | TCT | TCT | TTA | CGA | AAT | CTA | AAG | CTT | CTA | ATC | TAT | TTG |
| Heera | H | A | T | H | V | P | S | S | L | R | N | L | K | L | L | I | Y | L |
| | CAT | GCA | ACT | CAT | GTA | CCT | TCT | TCT | TTA | CGA | AAT | CTA | AAG | CTT | CTA | ATC | TAT | TTG |
| Skorospieka | H | A | T | H | V | P | S | S | L | R | N | L | K | L | L | I | Y | L |
| | CAT | GCA | ACT | CAT | GTA | CCT | TCT | TCT | TTA | CGA | AAT | CTA | AAG | CTT | CTA | ATC | TAT | TTG |
| Kranti | H | A | T | H | V | P | S | S | L | R | N | L | K | L | L | I | Y | L |
| | CAT | GCA | ACT | CAT | GTA | CCT | TCT | TCT | TTA | CGA | AAT | CTA | AAG | CTT | CTA | ATC | TAT | TTG |
| Tumida | H | A | T | H | V | P | S | S | L | R | N | L | K | L | L | I | Y | L |
| | CAT | GCA | ACT | CAT | GTA | CCT | TCT | TCT | TTA | CGA | AAT | CTA | AAG | CTT | CTA | ATC | TAT | TTG |

1900 1910 1920 1930 1940

| | | | | | | | | | | | | | | | | | | |
|---|---|---|---|---|---|---|---|---|---|---|---|---|---|---|---|---|---|---|
| Donskaja-IV | R | I | Y | S | W | K | R | V | H | L | P | S | I | F | K | E | M | V |
| | AGA | ATT | TAC | TCC | TGG | AAA | AGA | GTT | CAT | CTG | CCA | AGT | ATC | TTT | AAA | GAG | ATG | GTA |
| Cutlass | R | I | Y | S | W | K | R | V | H | L | P | S | I | F | K | E | M | V |
| | AGA | ATT | TAC | TCC | TGG | AAA | AGA | GTT | CAT | CTG | CCA | AGT | ATC | TTT | AAA | GAG | ATG | GTA |
| Varuna | S | I | Y | S | R | E | R | V | H | L | P | S | I | F | K | E | M | V |
| | AGT | ATT | TAC | TCC | CGG | GAA | AGA | GTT | CAT | CTG | CCA | AGT | ATC | TTT | AAA | GAG | ATG | GTA |
| Heera | S | I | Y | S | R | E | R | V | H | L | P | S | I | F | K | E | M | V |
| | AGT | ATT | TAC | TCC | CGG | GAA | AGA | GTT | CAT | CTG | CCA | AGT | ATC | TTT | AAA | GAG | ATG | GTA |
| Skorospieka | S | I | Y | S | R | E | R | V | H | L | P | S | I | F | K | E | M | V |
| | AGT | ATT | TAC | TCC | CGG | GAA | AGA | GTT | CAT | CTG | CCA | AGT | ATC | TTT | AAA | GAG | ATG | GTA |
| Kranti | S | I | Y | S | R | E | R | V | H | L | P | S | I | F | K | E | M | V |
| | AGT | ATT | TAC | TCC | CGG | GAA | AGA | GTT | CAT | CTG | CCA | AGT | ATC | TTT | AAA | GAG | ATG | GTA |
| Tumida | S | I | Y | S | R | E | R | V | H | L | P | S | I | F | K | E | M | V |
| | AGT | ATT | TAC | TCC | CGG | GAA | AGA | GTT | CAT | CTG | CCA | AGT | ATC | TTT | AAA | GAG | ATG | GTA |

1950 1960 1970 1980 1990

| | | | | | | | | | | | | | | | | | | |
|---|---|---|---|---|---|---|---|---|---|---|---|---|---|---|---|---|---|---|
| Donskaja-IV | E | L | R | F | L | I | L | P | R | S | F | D | A | K | T | K | L | E |
| | GAG | TTG | AGG | TTC | CTG | ATT | CTT | CCC | CGT | TCT | TTT | GAC | GCT | AAA | ACA | AAG | TTG | GAA |
| Cutlass | E | L | R | F | L | I | L | P | R | S | F | D | A | K | T | K | L | E |
| | GAG | TTG | AGG | TTC | CTG | ATT | CTT | CCC | CGT | TCT | TTT | GAC | GCT | AAA | ACA | AAG | TTG | GAA |
| Varuna | Q | L | R | F | L | V | L | P | P | F | V | D | A | R | T | K | L | E |
| | CAG | TTG | AGG | TTC | CTG | GTC | CTT | CCC | CCT | TTT | GTT | GAC | GCT | AGA | ACA | AAG | TTG | GAA |
| Heera | Q | L | R | F | L | V | L | P | P | F | V | D | A | R | T | K | L | E |
| | CAG | TTG | AGG | TTC | CTG | GTC | CTT | CCC | CCT | TTT | GTT | GAC | GCT | AGA | ACA | AAG | TTG | GAA |
| Skorospieka | Q | L | R | F | L | V | L | P | P | F | V | D | A | R | T | K | L | E |
| | CAG | TTG | AGG | TTC | CTG | GTC | CTT | CCC | CCT | TTT | GTT | GAC | GCT | AGA | ACA | AAG | TTG | GAA |
| Kranti | Q | L | R | F | L | V | L | P | P | F | V | D | A | R | T | K | L | E |
| | CAG | TTG | AGG | TTC | CTG | GTC | CTT | CCC | CCT | TTT | GTT | GAC | GCT | AGA | ACA | AAG | TTG | GAA |
| Tumida | Q | L | R | F | L | V | L | P | P | F | V | D | A | R | T | K | L | E |
| | CAG | TTG | AGG | TTC | CTG | GTC | CTT | CCC | CCT | TTT | GTT | GAC | GCT | AGA | ACA | AAG | TTG | GAA |

2000 2010 2020 2030 2040 2050

| | | | | | | | | | | | | | | | | | | |
|---|---|---|---|---|---|---|---|---|---|---|---|---|---|---|---|---|---|---|
| Donskaja-IV | L | G | N | L | V | N | L | E | C | L | T | G | F | R | S | E | Y | G |
| | TTG | GGT | AAT | CTA | GTG | AAC | CTG | GAG | TGC | TTG | ACC | GGC | TTC | CGA | TCA | GAA | TAC | GGT |
| Cutlass | L | G | N | L | V | N | L | E | C | L | T | G | F | R | S | E | Y | G |
| | TTG | GGT | AAT | CTA | GTG | AAC | CTG | GAG | TGC | TTG | ACC | GGC | TTC | CGA | TCA | GAA | TAC | GGT |
| Varuna | L | G | N | L | V | N | L | E | Y | L | I | C | F | Q | S | Q | Y | G |
| | CTG | GGT | AAT | CTA | GTG | AAC | CTG | GAG | TAC | TTG | ATC | TGC | TTC | CAA | TCA | CAA | TAC | GGT |
| Heera | L | G | N | L | V | N | L | E | Y | L | I | C | F | Q | S | Q | Y | G |
| | CTG | GGT | AAT | CTA | GTG | AAC | CTG | GAG | TAC | TTG | ATC | TGC | TTC | CAA | TCA | CAA | TAC | GGT |
| Skorospieka | L | G | N | L | V | N | L | E | Y | L | I | C | F | Q | S | Q | Y | G |
| | CTG | GGT | AAT | CTA | GTG | AAC | CTG | GAG | TAC | TTG | ATC | TGC | TTC | CAA | TCA | CAA | TAC | GGT |
| Kranti | L | G | N | L | V | N | L | E | Y | L | I | C | F | Q | S | Q | Y | G |
| | CTG | GGT | AAT | CTA | GTG | AAC | CTG | GAG | TAC | TTG | ATC | TGC | TTC | CAA | TCA | CAA | TAC | GGT |
| Tumida | L | G | N | L | V | N | L | E | Y | L | I | C | F | Q | S | Q | Y | G |
| | CTG | GGT | AAT | CTA | GTG | AAC | CTG | GAG | TAC | TTG | ATC | TGC | TTC | CAA | TCA | CAA | TAC | GGT |

2060 2070 2080 2090 2100

| | | | | | | | | | | | | | | | | | | |
|---|---|---|---|---|---|---|---|---|---|---|---|---|---|---|---|---|---|---|
| Donskaja-IV | S | I | T | D | F | L | R | M | K | K | L | R | T | L | E | I | F | L |
| | AGC | ATC | ACA | GAC | TTC | CTC | CGA | ATG | AAA | AAG | CTC | AGG | ACT | CTC | GAG | ATA | TTT | CTC |
| Cutlass | S | I | T | D | F | L | R | M | K | K | L | R | T | L | E | I | F | L |
| | AGC | ATC | ACA | GAC | TTC | CTC | CGA | ATG | AAA | AAG | CTC | AGG | ACT | CTC | GAG | ATA | TTT | CTC |
| Varuna | S | I | T | D | F | L | R | M | N | K | L | R | N | L | H | I | Q | L |
| | AGC | ATC | ACA | GAC | TTC | CTC | CGA | ATG | AAT | AAG | CTC | AGG | AAT | CTC | CAC | ATA | CAA | CTC |
| Heera | S | I | T | D | F | L | R | M | N | K | L | R | N | L | H | I | Q | L |
| | AGC | ATC | ACA | GAC | TTC | CTC | CGA | ATG | AAT | AAG | CTC | AGG | AAT | CTC | CAC | ATA | CAA | CTC |
| Skorospieka | S | I | T | D | F | L | R | M | N | K | L | R | N | L | H | I | Q | L |
| | AGC | ATC | ACA | GAC | TTC | CTC | CGA | ATG | AAT | AAG | CTC | AGG | AAT | CTC | CAC | ATA | CAA | CTC |
| Kranti | S | I | T | D | F | L | R | M | N | K | L | R | N | L | H | I | Q | L |
| | AGC | ATC | ACA | GAC | TTC | CTC | CGA | ATG | AAT | AAG | CTC | AGG | AAT | CTC | CAC | ATA | CAA | CTC |
| Tumida | S | I | T | D | F | L | R | M | N | K | L | R | N | L | H | I | Q | L |
| | AGC | ATC | ACA | GAC | TTC | CTC | CGA | ATG | AAT | AAG | CTC | AGG | AAT | CTC | CAC | ATA | CAA | CTC |

2110 2120 2130 2140 2150 2160

| | | | | | | | | | | | | | | | | | | |
|---|---|---|---|---|---|---|---|---|---|---|---|---|---|---|---|---|---|---|
| Donskaja-IV | K | G | R | Y | T | S | E | T | L | A | S | S | L | C | E | L | R | N |
| | AAA | GGC | AGG | TAT | ACT | TCT | GAA | ACT | CTA | GCG | TCA | TCT | CTC | TGT | GAA | TTA | AGA | AAC |
| Cutlass | K | G | - | Y | T | S | E | T | L | A | S | S | L | C | E | L | R | N |
| | AAA | GGG | --- | TAT | ACT | TCT | GAA | ACT | CTA | GCG | TCA | TCT | CTC | TGT | GAA | TTA | AGA | AAC |
| Varuna | K | G | - | Y | T | S | E | I | L | A | S | S | L | C | E | L | R | K |
| | AAA | GGG | --- | TAT | ACT | TCT | GAA | ATT | CTA | GCG | TCA | TCT | CTC | TGT | GAA | TTA | AGA | AAA |
| Heera | K | G | - | Y | T | S | E | I | L | A | S | S | L | C | E | L | R | K |
| | AAA | GGG | --- | TAT | ACT | TCT | GAA | ATT | CTA | GCG | TCA | TCT | CTC | TGT | GAA | TTA | AGA | AAA |
| Skorospieka | K | G | - | Y | T | S | E | I | L | A | S | S | L | C | E | L | R | K |
| | AAA | GGG | --- | TAT | ACT | TCT | GAA | ATT | CTA | GCG | TCA | TCT | CTC | TGT | GAA | TTA | AGA | AAA |
| Kranti | K | G | - | Y | T | S | E | I | L | A | S | S | L | C | E | L | R | K |
| | AAA | GGG | --- | TAT | ACT | TCT | GAA | ATT | CTA | GCG | TCA | TCT | CTC | TGT | GAA | TTA | AGA | AAA |
| Tumida | K | G | - | Y | T | S | E | I | L | A | S | S | L | C | E | L | R | K |
| | AAA | GGG | --- | TAT | ACT | TCT | GAA | ATT | CTA | GCG | TCA | TCT | CTC | TGT | GAA | TTA | AGA | AAA |

2170 2180 2190 2200 2210

| | | | | | | | | | | | | | | | | | | |
|---|---|---|---|---|---|---|---|---|---|---|---|---|---|---|---|---|---|---|
| Donskaja-IV | L | E | V | L | R | L | I | D | E | N | R | - | S | G | G | A | Y | D |
| | CTG | GAG | GTG | CTT | AGG | TTG | ATC | GAT | GAG | AAC | CGA | --- | TCC | GGT | GGG | GCT | TAT | GAT |
| Cutlass | L | E | V | L | R | L | I | D | E | N | N | G | S | G | G | A | Y | D |
| | CTG | GAG | GTG | CTT | AGG | TTG | ATC | GAT | GAG | AAC | AAC | GGA | TCC | GGT | GGG | GCT | TAT | GAT |
| Varuna | L | E | E | L | S | L | I | D | E | N | N | E | S | D | G | A | Y | D |
| | CTG | GAG | GAG | CTT | AGC | TTG | ATC | GAT | GAG | AAC | AAC | GAA | TCC | GAT | GGG | GCT | TAT | GAT |
| Heera | L | E | E | L | S | L | I | D | E | N | N | E | S | D | G | A | Y | D |
| | CTG | GAG | GAG | CTT | AGC | TTG | ATC | GAT | GAG | AAC | AAC | GAA | TCC | GAT | GGG | GCT | TAT | GAT |
| Skorospieka | L | E | E | L | S | L | I | D | E | N | N | E | S | D | G | A | Y | D |
| | CTG | GAG | GAG | CTT | AGC | TTG | ATC | GAT | GAG | AAC | AAC | GAA | TCC | GAT | GGG | GCT | TAT | GAT |
| Kranti | L | E | E | L | S | L | I | D | E | N | N | E | S | D | G | A | Y | D |
| | CTG | GAG | GAG | CTT | AGC | TTG | ATC | GAT | GAG | AAC | AAC | GAA | TCC | GAT | GGG | GCT | TAT | GAT |
| Tumida | L | E | E | L | S | L | I | D | E | N | N | E | S | D | G | A | Y | D |
| | CTG | GAG | GAG | CTT | AGC | TTG | ATC | GAT | GAG | AAC | AAC | GAA | TCC | GAT | GGG | GCT | TAT | GAT |

2220 2230 2240 2250 2260

| | | | | | | | | | | | | | | | | | | |
|---|---|---|---|---|---|---|---|---|---|---|---|---|---|---|---|---|---|---|
| Donskaja-IV | V | D | F | V | W | N | F | I | H | L | R | S | L | K | L | G | I | R |
| | GTG | GAT | TTC | GTT | TGG | AAT | TTC | ATT | CAT | CTA | AGG | TCT | TTA | AAG | CTG | GGA | ATA | CGT |
| Cutlass | V | D | F | V | W | N | F | I | H | L | R | S | L | K | L | G | I | R |
| | GTG | GAT | TTC | GTT | TGG | AAT | TTC | ATT | CAT | CTA | AGG | TCT | TTA | AAG | CTG | GGA | ATA | CGT |
| Varuna | V | D | F | V | W | N | F | I | H | L | R | R | L | G | L | S | I | H |
| | GTG | GAT | TTC | GTT | TGG | AAT | TTC | ATT | CAT | CTA | AGG | CGT | TTA | GGG | CTG | AGC | ATA | CAT |
| Heera | V | D | F | V | W | N | F | I | H | L | R | R | L | G | L | S | I | H |
| | GTG | GAT | TTC | GTT | TGG | AAT | TTC | ATT | CAT | CTA | AGG | CGT | TTA | GGG | CTG | AGC | ATA | CAT |
| Skorospieka | V | D | F | V | W | N | F | I | H | L | R | R | L | G | L | S | I | H |
| | GTG | GAT | TTC | GTT | TGG | AAT | TTC | ATT | CAT | CTA | AGG | CGT | TTA | GGG | CTG | AGC | ATA | CAT |
| Kranti | V | D | F | V | W | N | F | I | H | L | R | R | L | G | L | S | I | H |
| | GTG | GAT | TTC | GTT | TGG | AAT | TTC | ATT | CAT | CTA | AGG | CGT | TTA | GGG | CTG | AGC | ATA | CAT |
| Tumida | V | D | F | V | W | N | F | I | H | L | R | R | L | G | L | S | I | H |
| | GTG | GAT | TTC | GTT | TGG | AAT | TTC | ATT | CAT | CTA | AGG | CGT | TTA | GGG | CTG | AGC | ATA | CAT |

2270 2280 2290 2300 2310

Donskaja-IV
I K K L P E H S R F P P H L A H V S
ATC AAA AAG CTT CCT GAG CAC TCT CGA TTT CCT CCC CAC CTT GCA CAT GTA TCT

Cutlass
I K K L P E H S R F P P H L A H V S
ATC AAA AAG CTT CCT GAG CAC TCT CGA TTT CCT CCC CAC CTT GCA CAT GTA TCT

Varuna
G T R L P D H S R F P P H L A H I T
GGG ACA AGG CTT CCT GAT CAC TCT CGA TTT CCT CCC CAC CTT GCA CAC ATA ACT

Heera
G T R L P D H S R F P P H L A H I T
GGG ACA AGG CTT CCT GAT CAC TCT CGA TTT CCT CCC CAC CTT GCA CAC ATA ACT

Skorospieka
G T R L P D H S R F P P H L A H I T
GGG ACA AGG CTT CCT GAT CAC TCT CGA TTT CCT CCC CAC CTT GCA CAC ATA ACT

Kranti
G T R L P D H S R F P P H L A H I T
GGG ACA AGG CTT CCT GAT CAC TCT CGA TTT CCT CCC CAC CTT GCA CAC ATA ACT

Tumida
G T R L P D H S R F P P H L A H I T
GGG ACA AGG CTT CCT GAT CAC TCT CGA TTT CCT CCC CAC CTT GCA CAC ATA ACT

2320 2330 2340 2350 2360 2370

Donskaja-IV
L S G C K M E E D P L Q I L E K L L
CTA AGT GGC TGT AAA ATG GAG GAG GAT CCA CTG CAG ATT CTG GAA AAG TTG CTT

Cutlass
L S G C K M E E D P L Q I L E K L L
CTA AGT GGC TGT AAA ATG GAG GAG GAT CCA CTG CAG ATT CTG GAA AAG TTG CTT

Varuna
L R D C E M E D D P L Q I L E K L L
CTA CGT GAC TGT GAA ATG GAG GAT GAT CCA CTG CAG ATT CTG GAA AAG TTG CTT

Heera
L R D C E M E D D P L Q I L E K L L
CTA CGT GAC TGT GAA ATG GAG GAT GAT CCA CTG CAG ATT CTG GAA AAG TTG CTT

Skorospieka
L R D C E M E D D P L Q I L E K L L
CTA CGT GAC TGT GAA ATG GAG GAT GAT CCA CTG CAG ATT CTG GAA AAG TTG CTT

Kranti
L R D C E M E D D P L Q I L E K L L
CTA CGT GAC TGT GAA ATG GAG GAT GAT CCA CTG CAG ATT CTG GAA AAG TTG CTT

Tumida
L R D C E M E D D P L Q I L E K L L
CTA CGT GAC TGT GAA ATG GAG GAT GAT CCA CTG CAG ATT CTG GAA AAG TTG CTT

2380 2390 2400 2410 2420

Donskaja-IV
H L K S V L L G I D A F V G R K M V
CAT TTA AAG TCA GTT TTA TTA GGA ATT GAT GCT TTC GTG GGG AGG AAG ATG GTG

Cutlass
H L K S V L L N Q .
CAT TTA AAG TCA GTT TTA TTA AAT CAA TGA

Varuna
H L K S A V S H Q R S F V G R K M V
CAT TTG AAG TCG GCT GTA TCA CAT CAA AGA TCT TTC GTG GGG AGG AAG ATG GTG

Heera
H L K S A V S H Q R S F V G R K M V
CAT TTG AAG TCG GCT GTA TCA CAT CAA AGA TCT TTC GTG GGG AGG AAG ATG GTG

Skorospieka
H L K S A V S H Q R S F V G R K M V
CAT TTG AAG TCG GCT GTA TCA CAT CAA AGA TCT TTC GTG GGG AGG AAG ATG GTG

Kranti
H L K S A V S H Q R S F V G R K M V
CAT TTG AAG TCG GCT GTA TCA CAT CAA AGA TCT TTC GTG GGG AGG AAG ATG GTG

Tumida
H L K S A V S H Q R S F V G R K M V
CAT TTG AAG TCG GCT GTA TCA CAT CAA AGA TCT TTC GTG GGG AGG AAG ATG GTG

2430 2440 2450 2460 2470 2480

Donskaja-IV
C S K G G F P Q L C K L L I W S L D
TGT TCA AAA GGT GGA TTC CCT CAA TTA TGT AAG CTA CTA ATA TGG TCC CTA GAC

Cutlass

Varuna
C S K G G F P Q L C K L Q I E L L D
TGT TCA AAA GGT GGA TTC CCT CAA TTA TGT AAG CTA CAA ATA GAG TTA CTA GAC

Heera
C S K G G F P Q L C K L Q I E L L D
TGT TCA AAA GGT GGA TTC CCT CAA TTA TGT AAG CTA CAA ATA GAG TTA CTA GAC

Skorospieka
C S K G G F P Q L C K L Q I E L L D
TGT TCA AAA GGT GGA TTC CCT CAA TTA TGT AAG CTA CAA ATA GAG TTA CTA GAC

Kranti
C S K G G F P Q L C K L Q I E L L D
TGT TCA AAA GGT GGA TTC CCT CAA TTA TGT AAG CTA CAA ATA GAG TTA CTA GAC

Tumida
C S K G G F P Q L C K L Q I E L L D
TGT TCA AAA GGT GGA TTC CCT CAA TTA TGT AAG CTA CAA ATA GAG TTA CTA GAC

2490 2500 2510 2520 2530

Donskaja-IV
D W E E W I V E E G S M P C L R T L
GAC TGG GAA GAG TGG ATA GTC GAA GAA GGG TCG ATG CCA TGT CTT CGT ACC TTG

Cutlass

Varuna
D W E E W I V E E G S M P C L R T L
GAC TGG GAA GAG TGG ATA GTC GAA GAA GGG TCG ATG CCA TGT CTT CGT ACC TTG

Heera
D W E E W I V E E G S M P C L R T L
GAC TGG GAA GAG TGG ATA GTC GAA GAA GGG TCG ATG CCA TGT CTT CGT ACC TTG

Skorospieka
D W E E W I V E E G S M P C L R T L
GAC TGG GAA GAG TGG ATA GTC GAA GAA GGG TCG ATG CCA TGT CTT CGT ACC TTG

Kranti
D W E E W I V E E G S M P C L R T L
GAC TGG GAA GAG TGG ATA GTC GAA GAA GGG TCG ATG CCA TGT CTT CGT ACC TTG

Tumida
D W E E W I V E E G S M P C L R T L
GAC TGG GAA GAG TGG ATA GTC GAA GAA GGG TCG ATG CCA TGT CTT CGT ACC TTG

2540 2550 2560 2570 2580

Donskaja-IV
S I W L C D K L K E L P E G L K Y I
TCT ATA TGG TTA TGT GAC AAG TTG AAG GAG CTA CCG GAA GGG CTC AAG TAT ATC

Cutlass

Varuna
S I W Y C E K L K E L P E G L K Y I
TCT ATA TGG TAT TGT GAA AAG TTG AAG GAG CTA CCG GAA GGG CTC AAG TAT ATC

Heera
S I W Y C E K L K E L P E G L K Y I
TCT ATA TGG TAT TGT GAA AAG TTG AAG GAG CTA CCG GAA GGG CTC AAG TAT ATC

Skorospieka
S I W Y C E K L K E L P E G L K Y I
TCT ATA TGG TAT TGT GAA AAG TTG AAG GAG CTA CCG GAA GGG CTC AAG TAT ATC

Kranti
S I W Y C E K L K E L P E G L K Y I
TCT ATA TGG TAT TGT GAA AAG TTG AAG GAG CTA CCG GAA GGG CTC AAG TAT ATC

Tumida
S I W Y C E K L K E L P E G L K Y I
TCT ATA TGG TAT TGT GAA AAG TTG AAG GAG CTA CCG GAA GGG CTC AAG TAT ATC

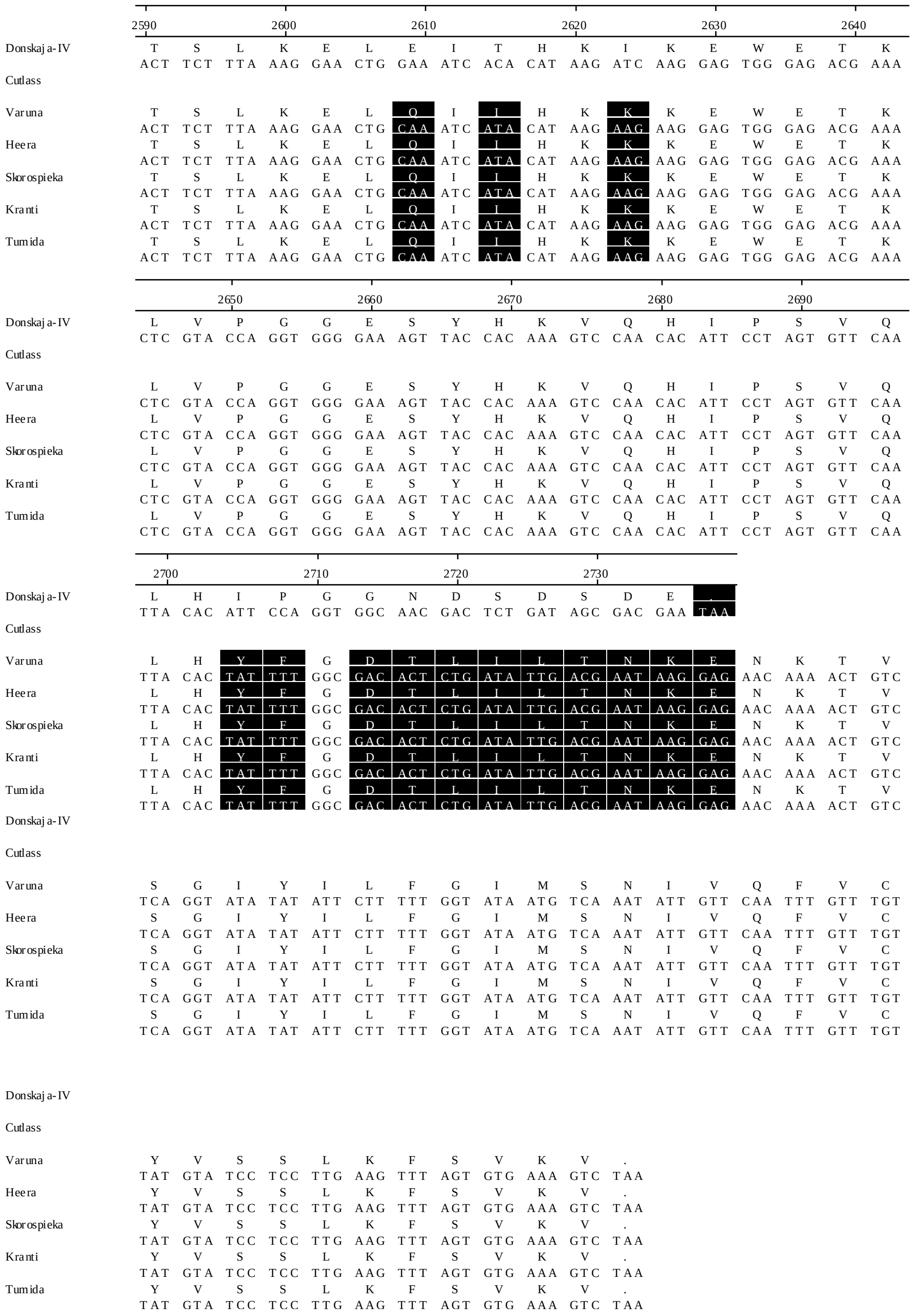


**Fig. S3.** Alignment of the nucleotide sequences of the coding region of *BjuWRR1* alleles and the amino acid sequences of the encoded proteins. Non-synonymous substitutions in different alleles with Donskaja-IV as the reference sequence, are highlighted in black.
